# Reflective multi-immersion microscope objectives inspired by the Schmidt telescope

**DOI:** 10.1101/2022.10.13.511906

**Authors:** Fabian F. Voigt, Anna Maria Reuss, Thomas Naert, Sven Hildebrand, Martina Schaettin, Adriana L. Hotz, Lachlan Whitehead, Armin Bahl, Stephan C. F. Neuhauss, Alard Roebroeck, Esther T. Stoeckli, Soeren S. Lienkamp, Adriano Aguzzi, Fritjof Helmchen

## Abstract

Rapid advances in tissue clearing protocols have begun to outpace the capabilities of existing microscope objectives: High-resolution imaging inside cm-sized cleared samples is often not possible as it requires multi-immersion objectives with high numerical aperture (NA > 0.7), long working distance (WD > 10 mm) and a large field-of-view (FOV > 1 mm). Here, we introduce a novel mirror-based optical design, the “Schmidt objective”, which meets all these criteria despite containing only two optical elements. It consists of a spherical mirror in contact with the immersion medium and an aspherical correction plate. We showcase a multi-photon variant of a Schmidt objective that reaches NA 1.08 at an refractive index of 1.56 and demonstrate its versatility by imaging fixed samples in a wide range of immersion media ranging from air and water to BABB, DBE, and ECI. In addition, we demonstrate in vivo imaging by recording neuronal activity in larval zebrafish.

## Introduction

Imaging is crucial for biology and microscope objectives thus are key tools for discovery. Since the pioneering work by Ernst Abbe 150 years ago, many engineering innovations have drastically improved their performance^1^. Nonetheless, application domains exist, for which better and more cost-efficient objectives still are highly desirable. One of these domains is imaging samples processed with tissue clearing techniques^2^ which render entire organs or organisms transparent while retaining fluorescent labeling. Given that over the course of the past decade, clearing of entire mouse brains (**≈** 1 cm^3^) has become routine^2–4^ and that clearing of entire mice and human organs is feasible^5–7^, there is a need for microscope objectives that are suitable for imaging large cleared samples at high resolution. This demand poses a challenging optical engineering problem: In microscopy, imaging at high resolution requires increasing the NA, which complicates the correction of aberrations and necessitates more lens elements. Beyond that, the large size of cleared samples requires objectives with long working distances while at the same time large FOVs are necessary to allow imaging with high throughput. The most challenging engineering problem by far is, however, that published clearing protocols require immersion media with a diversity of the index of refraction (n), ranging from water (n=1.33) for expansion microscopy^8^ to typical organic solvents such as dibenzyl ether^9^ (DBE) or ethyl cinnamate^10^ (ECI) with n=1.56. Some commercial objectives have a correction collar that moves an internal lens group to minimize index-dependent aberrations, but for high-NA objectives, designers often opt for narrowing the index range to keep the costs under control. For example, the Olympus XLSLPLN25XGMP combines NA=1.0 with 8-mm WD, but is restricted to n = 1.41 - 1.52. Covering a larger index range is possible at the expense of resolution (e.g. the Olympus XLPLN10XSVMP has NA 0.6, 8-mm WD, and n=1.33 - 1.52). Such objectives typically have 13-15 lenses and are expensive (**≈** 25-30 k$). In addition, despite this considerable effort, indices around 1.56 cannot be reached with satisfactory image quality. And not least, media such as DBE, ECI, or benzyl alcohol / benzyl benzoate (BABB) can dissolve lens cements so that chemically resistant dipping cap designs are required. Few such options are available, of which one is an ASI/Special Optics objective (54-12-8) that covers the entire range from 1.33 to 1.56 with good chemical compatibility, featuring 10-mm WD at NA 0.7. However, scaling this design up to higher NA, longer WD, and a FOV beyond 1 mm has not been achieved so far. To address this challenge, we introduce a novel approach for designing a multi-immersion objective based on a mirror instead of lenses. The underlying idea is that the reflection of a light ray by a mirror is independent of the index of refraction of the medium the mirror is in contact with. Consequently, if we submerge a curved mirror inside a liquid-filled chamber and use it to focus light, the ray paths and location of the focus do not change when changing the bulk index of the immersion medium. An invariant ray path means that any monochromatic aberrations such as spherical aberration or coma stay constant even though the numerical aperture of the objective scales with the refractive index of the medium. This is drastically different compared to lens-based multi-immersion designs where any medium change causes additional aberrations that need to be accounted for. In addition, reflective objective designs often have a longer WD than refractive designs at comparable NA. This combination of features means that reflective optical systems are excellent templates for long-WD multi-immersion objectives. Although to our knowledge such design has not been previously applied in bioimaging, it is found in nature, where each of the hundreds of eyes of scallops contains a curved mirror to form an image^11^ (Figure 1a). Each mirror is in direct contact with the liquid-filled space containing the photoreceptors. Each eye also contains a lens, which is not, however, the primary image-forming element^12^. This combination of a lens and a mirror is reminiscent of the optical design of the Schmidt telescope^13^, a mirror-based wide-field telescope design commonly used in astronomy since the 1930s (Figure 1b). A Schmidt telescope is based on a spherical mirror which as a standalone element would create a heavily aberrated image due to spherical aberration. By adding a refractive aspherical correction plate in the center of curvature of the mirror, Bernhard Schmidt (1879-1935) created a telescope design that corrects the spherical aberration and provides excellent image quality over a large FOV. Owing to this wide-field capability, such telescopes are exceptionally well suited for surveys of the night sky. For example, the Kepler space telescope is a Schmidt telescope optimized for exoplanet detection^14,15.^ Most Schmidt telescopes share the characteristic that due to the rotational symmetry of the spherical mirror, the image surface of a Schmidt telescope is curved as well. The usage of an aspherical correction plate in combination with curved mirrors forms a general solution that can be applied to a wide variety of optical design problems. Indeed, shortly after the Schmidt telescope became widely known, it was recognized that this principle could be useful to build microscope objectives^17,18.^ However, apart from non-imaging flow cytometry^16^, there is no application of the Schmidt principle in modern biophotonics.

**Fig. 1.**
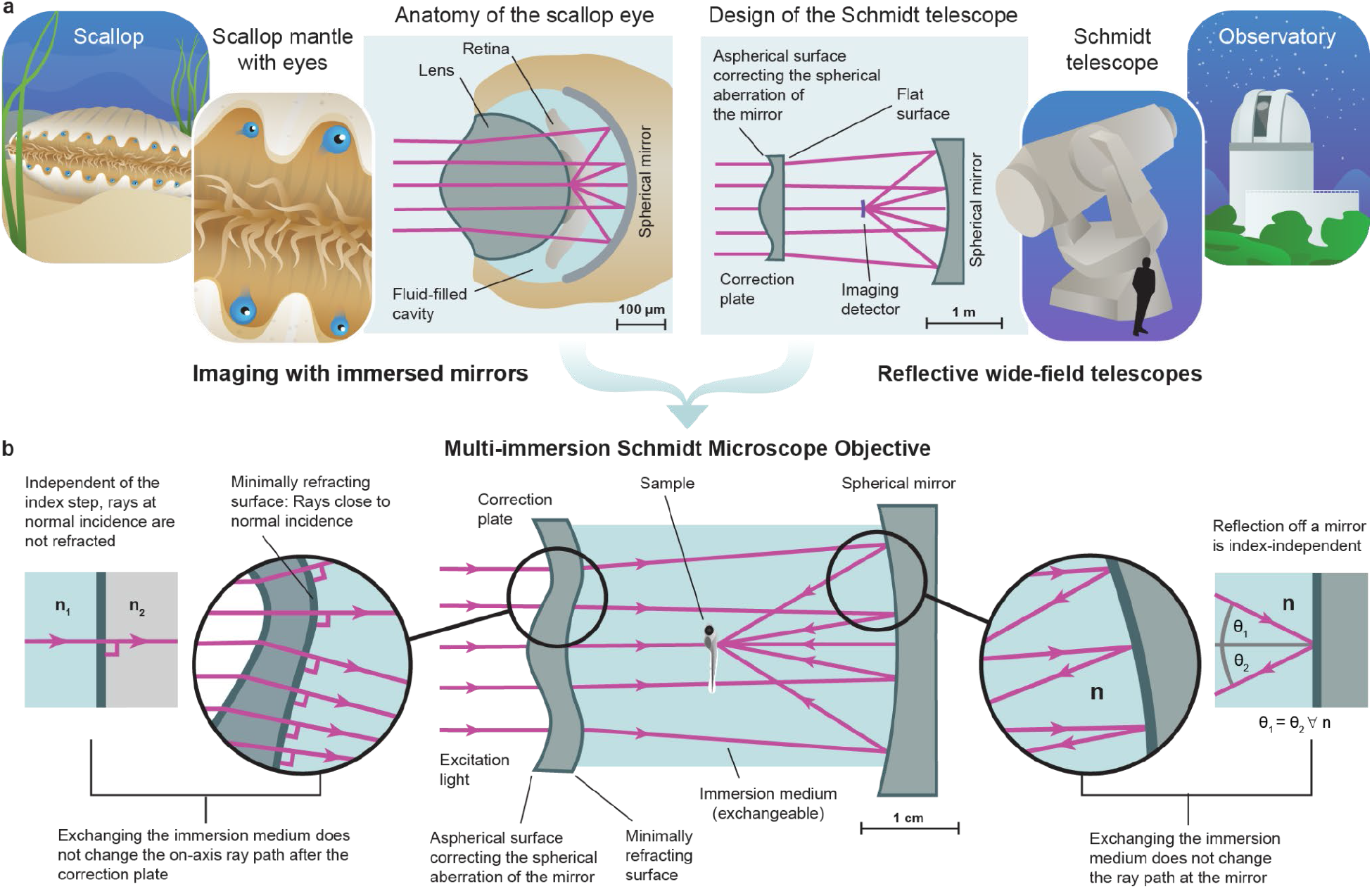
Concept of the multi-immersion Schmidt Objective. **a**, Our approach is inspired by the anatomy of the scallop eye: In this eye design, a curved mirror in contact with a liquid forms an image on a transparent layer of photoreceptors. A second inspiration is the Schmidt telescope, a mirror-based wide-field telescope design that consists of a spherical mirror and an aspherical correction plate. **b**, We synthesize both concepts into a multi-immersion objective design that provides a sharp image in any homogeneous medium (center). Reflection off the mirror does not depend on the refractive index n of the immersion medium (right). To avoid additional refraction at the inner surface of the correction plate, we deform this surface such that all passing rays are close to normal incidence (bottom left).

## Results

### Turning a Schmidt telescope into an immersion objective

We reasoned that it is possible to turn a Schmidt telescope into an immersion microscope objective by a series of conceptual steps (Figure 1c; Supplementary Video 1):

1. Downscaling the design from the meter-sized apertures common in astronomy to a cm-sized aperture suitable for a microscope;
2. Replacing the detector in the focus of the telescope with a fluorescently labeled sample; Focusing and scanning an excitation beam, e.g., a near-infrared ultrafast pulsed laser beam for two-photon microscopy, instead of focusing light from stars;
3. Filling the entire inner space between the mirror and the correction plate with an immersion liquid.

We noted that this approach results in objective designs with excellent simulated image quality, however, the optimal shape of the aspherical surface correcting for spherical aberration needs to be tailored specifically to each immersion medium. Such designs with simple Schmidt correction plates thus are immersion objectives, but not multi-immersion-capable. The reason is that refraction at the interface between correction plate and immersion medium introduces additional index-dependent spherical aberration. To circumvent this problem, we make use of an edge case of the law of refraction: No matter how big the index difference at a refractive interface is, a ray at normal incidence is not refracted, i.e., it does not change its propagation direction. Therefore, we additionally shape the inner surface of the correction plate such that at all locations the passing rays are at normal incidence. In other words, the first aspherical surface of the correction plate deforms the incoming parallel wavefront to counteract the spherical aberration of the spherical mirror and the second surface is then shaped exactly like this deformed wavefront. As the wavefront passes the interface unchanged, no additional (index-dependent) aberrations are introduced. We refer to this interface as a “minimally refractive” surface because it does not have any net refractive power. Because in typical laser scanning microscopes the scan angles at the objective are usually only a few degrees, even off-axis ray bundles strike the minimally refractive surface at near normal incidence, which allows the design to have mm-sized FOVs. We term the resulting multi-immersion Schmidt-telescope-turned-microscope-objective a “Schmidt objective”.

### Design of a multi-photon Schmidt objective

Whereas our concept can be used to design objectives for a wide variety of imaging modalities, we opted to demonstrate the design as a multi-photon objective. The reason is that many multi-photon techniques such as two-photon microscopy require only a single laser source and the excitation spectrum is so narrow that no color correction is necessary. Consequently, the design can be kept extremely simple: Our multi-photon objective design has only two optical elements and can be used with a tunable excitation laser in the 750-1000 nm range. Owing to the reflective geometry, the sample sits in between the mirror and correction plate. Therefore, we define the working distance as the axial spacing between the focus and the outer rim of the mirror because this is the feasible z-range before a sample holder would collide with the mirror (Extended Data Figure 1a). For our Schmidt objective this working distance is 11 mm.

As the on-axis ray path is independent of the refractive index of the immersion medium, the NA scales linearly with n and no correction collar is required. In air and using 800-nm excitation light, the design has an NA of 0.69 and a diffraction-limited field of view (dFOV) of 1.6 mm diameter (Extended Data Figure 1b). The system has an index-independent entrance pupil diameter of 22 mm. As a result, the etendue (or optical invariant of the system) is constant and the product of NA and FOV is invariant as well. Consequently, in a higher-index medium such as water, the NA increases to 0.92 but the dFOV decreases to 1.4 mm. For typical organic solvents such as ECI, DBE, or BABB with an index of 1.56, the NA reaches 1.08 and the dFOV is 1.1 mm. The focal surface of the Schmidt objective is curved with a radius of 15.68 mm. Supplementary Note 1 provides an overview of the theory underlying our design approach.

To turn this design into a prototype, we designed an immersion chamber around the optical components of the objective and integrated it into a custom horizontal multi-photon microscope (Figure 2a-c). We chose a horizontal setup as it makes the exchange of immersion media and samples straightforward. In our prototype, the mirror is attached to a motorized XYZ stage and needs to be aligned relative to the correction plate. We opted for this approach because initially it was unclear how often the mirror would require cleaning and possibly replacement due to damaged coatings. In practice, however, we did not observe any coating damage even after prolonged immersion of the mirror in imaging media such as DBE, BABB or ECI (Supplementary Note 1). This means that future Schmidt objectives can be designed with a mirror that is fixed in the immersion chamber.

**Fig. 2.**
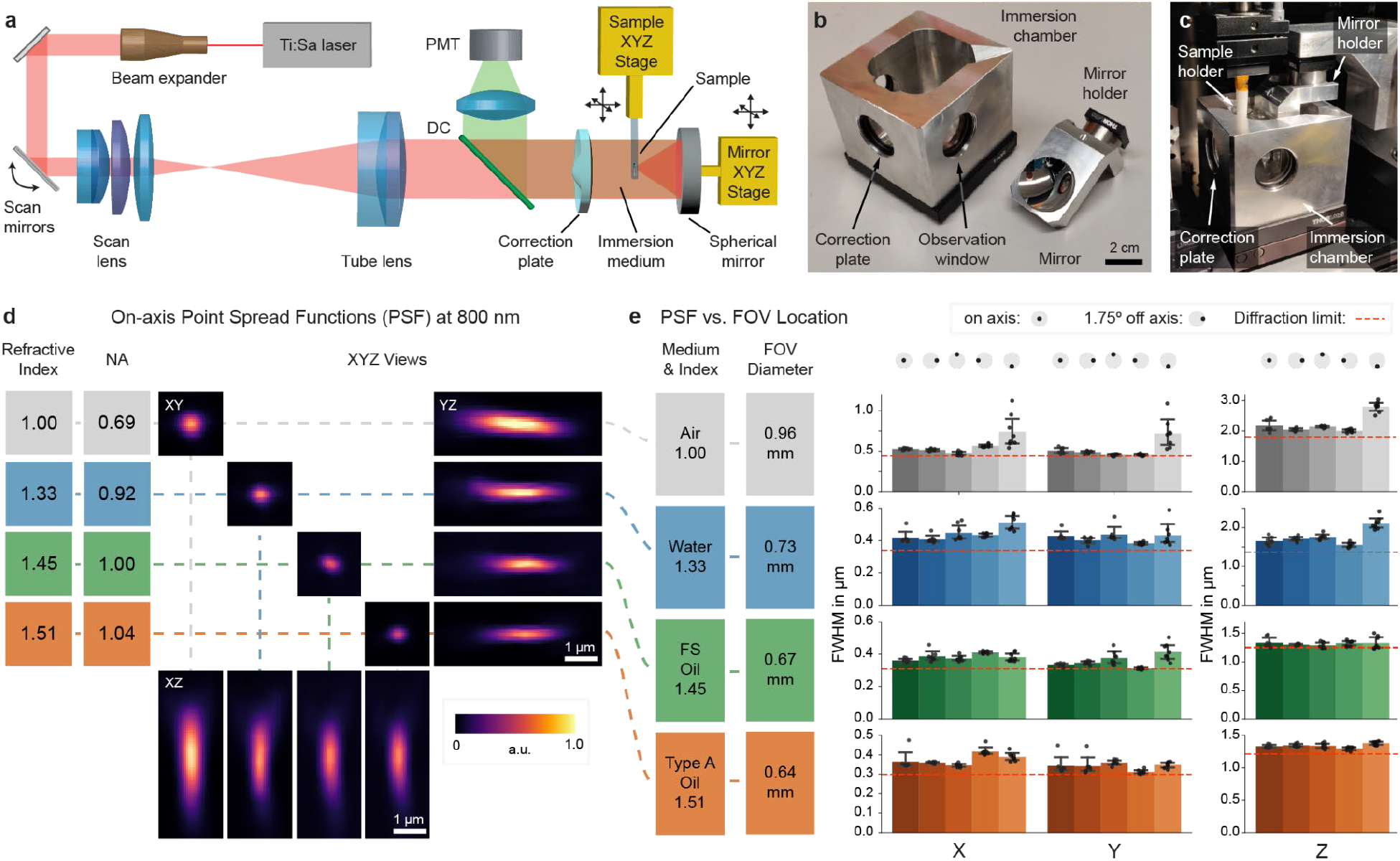
Setup and characterization. **a**, Overview of the setup. In our prototype, the position of the spherical mirror needs to be aligned relative to the correction plate for optimum image quality. This design allows for straightforward cleaning of the mirror when switching immersion media. To create a z-stack, the sample is moved along the optical axis. **b**, The prototype objective consists of an immersion chamber and a mirror. **c**, Assembled microscope objective. **d**, XYZ views of point spread function (PSF) measurements of 200-nm fluorescent beads imaged at refractive indices ranging from n=1.00 (air), n=1.33 (water), n=1.45 (Cargille fused silica matching oil), to n=1.51 (Cargille oil type A). Because NA) is proportional to n, an increase in n leads to higher NA and thus a smaller PSF. **e**, PSF uniformity over the field of view: PSF measurements were carried out on axis and at a scan angle of 1.75°, which was simulated to be the maximum theoretical diffraction-limited scan angle for the total system (Extended Data Figure 1). As the magnification of the objective is proportional to n, the FOV diameter decreases with increasing index. Diffraction-limited theoretical values are indicated by the red dashed lines.

### Characterization of the multi-photon Schmidt objective

We characterized the optical resolution of the objective by imaging 200-nm fluorescent beads at 800-nm excitation wavelength (Figure 2d,e). In air (NA=0.69), we measured an on-axis point-spread-function (PSF) with a full-width-at-half-maximum (FWHM) of 0.56 ± 0.01 μm in X (mean ± s.e.m.), 0.49 ± 0.07 μm in Y, and 1.93 ± 0.01 μm in Z (n=8 beads). In water, the PSF size decreased to 0.42 ± 0.04 μm in X, 0.43 ± 0.03 μm in Y and 1.65 ± 0.1 μm in Z (n=8 beads). When filling the objective with an immersion oil with n=1.45, the PSF size further decreased to 0.36 ± 0.01 μm in X, 0.33 ± 0.01 μm in Y, and 1.33 ± 0.09 μm in Z (n=8 beads). Further increasing the index to n=1.51 led to PSF size of 0.36 ± 0.02 μm in X, 0.32 ± 0.02 μm in Y, and 1.36 ± 0.01 μm in Z (n=8 beads). Across the index range, the on-axis PSF sizes were comparable to diffraction-limited values^9^. Imaging parameters are listed in Supplementary Table 1. According to our simulations (Extended Data Figure 1b-d), our objective supports a diffraction-limited FOV of 1.7 mm in air, 1.4 mm in water, and 1.1 mm at n=1.56. Due to additional off-axis aberrations introduced by our scan and tube lens, the whole system is diffraction-limited only up to a maximum scan angle of ±1.75° which corresponds to a dFOV of 0.74 mm in air, 0.68 mm in water, and 0.54 mm at n=1.56 (Extended Data Figure 1e-g). Therefore, we characterized the PSFs at a scan angle of ±1.75° (Figure 2e and Extended Data Figure 2) and found that the off-axis PSFs are in good agreement with our simulations.

### Imaging examples using the Schmidt objective

To demonstrate the imaging capabilities of our Schmidt objective, we imaged both cleared and living samples (Figure 3). In cleared samples, we first acquired low-resolution overview datasets with a mesoSPIM^10^ single-photon light-sheet microscope before transferring the sample into the Schmidt objective for high-resolution imaging. To showcase that the objective can be used in air, we imaged a pollen pellet composed of many individual pollen grains (Figure 3a). At higher zoom levels, surface features on individual pollen grains become readily apparent. Because the Schmidt objective is a multi-immersion objective, we could then fill the immersion chamber with ECI (n=1.56) and image the same pollen grains in this high-index medium (Extended Data Figure 3).

**Fig. 3.**
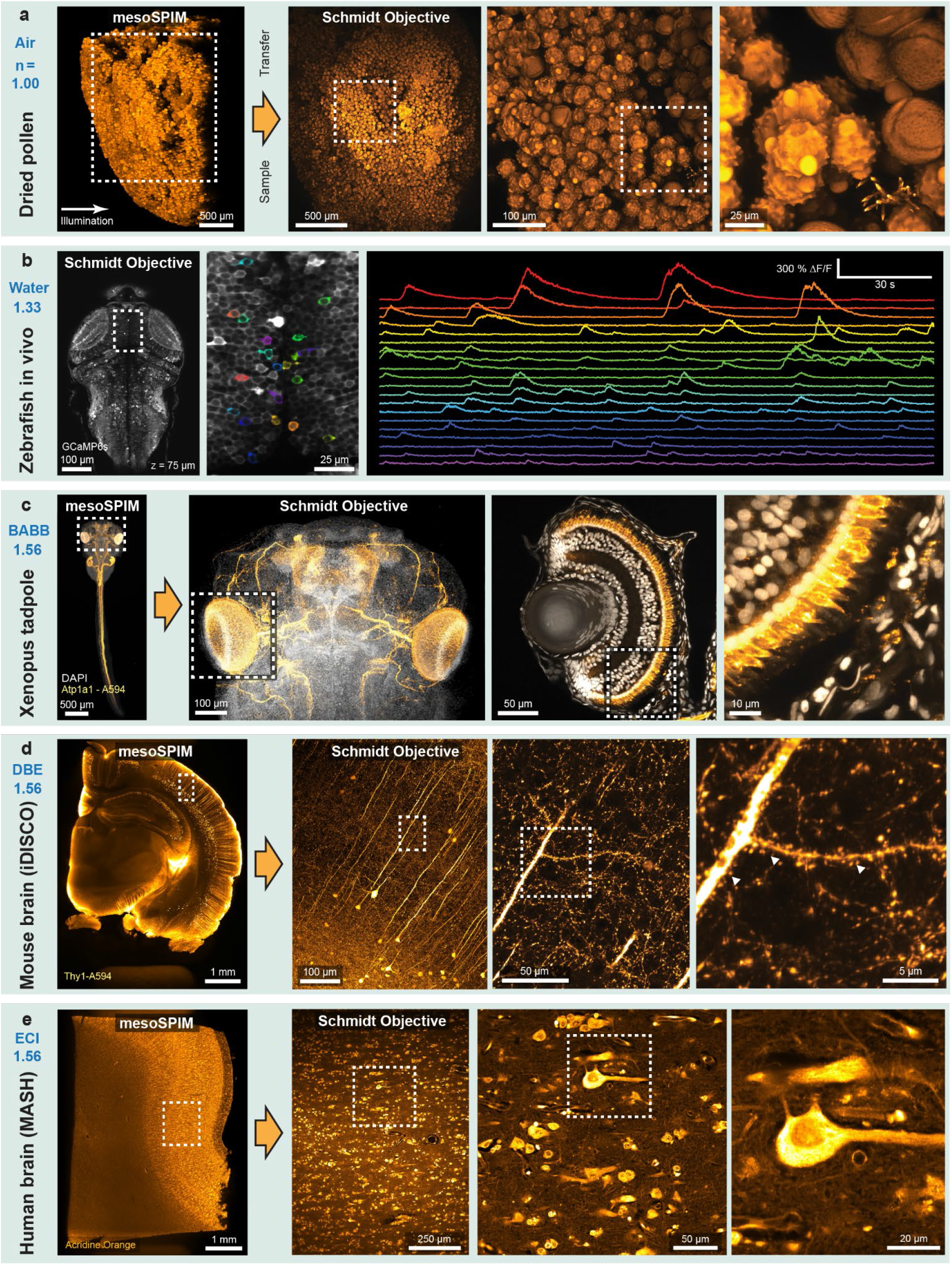
Example two-photon datasets acquired with the Schmidt objective. Fixed samples were first imaged with a mesoSPIM light-sheet microscope before the sample was transferred to the Schmidt objective. **a**, A pollen pellet composed of many individual pollen grains imaged in air. **b**, Functional imaging in a 5-day old elavl3:GCaMP6s zebrafish larvae in vivo. **c**, BABB-cleared *Xenopus tropicalis* tadpole stained for Atp1a1 (Alexa 594, orange) and nuclei (DAPI, greyscale). The large Schmidt FOV allows both imaging of the entire head (≈800 μm across) and imaging of individual developing photoreceptors in the eye (right). **d**, Thy1-H-labeled coronal mouse brain slice imaged in DBE. At higher magnification, spines on apical dendrites of L5 neurons are visible. **e**, MASH-processed human neocortex stained with Acridine Orange.

Next, we placed a 5-day old larval zebrafish expressing the calcium indicator GCaMP6s in neurons in the Schmidt objective and filled it with fish water (n=1.33). We acquired stacks of the entire zebrafish brain and were able to record neuronal activity in the tectum (Figure 3b). To illustrate that the objective can be used for imaging cleared samples, we imaged *Xenopus tropicalis* tadpoles in BABB (n=1.56), visualizing their developing peripheral nervous system and their eyes (Figure 3c and Supplementary Video 2). It was also possible to perform second-harmonic generation (SHG) imaging of muscle fibers in a tadpole tail, resolving individual sarcomeres (Extended Data Figure 4). We also applied the Schmidt objective to image larger cleared embryos in BABB, for example a 4-day old chicken embryo, which is about 6 mm in size (Extended Data Figure 5 & Supplementary Video 3). Next, we replaced the immersion medium with DBE (also n=1.56) and inserted a 4-mm thick coronal slice of a iDISCO-processed^3^ Thy1-YFP-H mouse brain. In this sample, we were able to visualize spines along apical dendrites (Figure 3d). We also successfully imaged dopaminergic neurons in a sample from a mouse brain stained for Tyrosine Hydroxylase (Extended Data Figure 6). Finally, we acquired a dataset from a MASH-processed^19^ 5×7×4 mm^3^ piece of human neocortex in ECI, enabling us to visualize neuronal somata and the dense neuropil (Figure 3e).

## Discussion

Taken together, these examples illustrate how a single Schmidt objective, due to its extreme index range, covers a range of applications that would usually require several objectives. Most importantly, *despite* having only three optical surfaces, our design outperforms all existing commercial multi-photon microscope objectives for cleared tissue within the parameter space of WD, index range, maximum NA, and FOV (Extended Data Figure 7). Whereas our mirror-based design provides multi-immersion capability basically for free, the reflective ray path means that the objective surrounds the sample which is similar to light-sheet microscopes for which custom sample holders and mounting techniques are often necessary. In addition, the bigger the sample is, the more it obstructs the excitation path. In principle, it is possible to design unobstructed off-axis Schmidt objectives to ameliorate these disadvantages. Similar to a Schmidt telescope, we allow the focal surface to be curved, which did not turn out to be an issue when imaging three-dimensional samples because the z-shift across the FOV is small (i.e. Δz = 1.2 μm across a 400 μm FOV).

A more important concern is that the long path length inside the immersion medium means that the image quality depends on the homogeneity of the liquid. For us, this meant that whereas imaging in organic solvents was straightforward, we could not achieve satisfactory image quality in CLARITY samples because the required refractive index matching solutions were not homogeneous enough. This property is not specific to the Schmidt objective, however: Any immersion objective with long WD and high NA will show similar behavior. We foresee that the ongoing development of clearing techniques for larger and larger samples will naturally converge towards highly homogeneous imaging media as they provide the best imaging results independent of the microscopy modality. We believe that our work opens up a previously unexplored design space for building immersion microscope objectives. In principle, our concept can be extended to any imaging modality, including widefield, confocal, and light-sheet microscopy. The underlying simplicity of the concept makes it well suited for low-cost objectives that can be mass-produced for diagnostic applications. Additionally, in optical design, simplicity begets scalability: Our approach will likely allow the design of “mesolenses” - objectives that perform high-resolution (NA>1) imaging across cm-scale FOVs and are capable of extremely high-throughput imaging^20^. Beyond that, our development of the Schmidt objective shows that astronomers and microscopists might benefit from closer interactions. In this regard, our work lends credence to a remark that Victor Hugo made in *Les Misérables* in 1862: “*Where the telescope ends the microscope begins, and who can say which of the two provides the grander view? Choose*.”

## Supporting information

Supplemental Video 1 - Overview

Supplemental Video 2 - Imaging a Xenopus tadpole

Supplemental Video 3 - Imaging a chicken embryo

## Acknowledgements

We would like to thank Stefan Giger for machining all custom objective parts, Julia Kuhl for illustrations, David Liittschwager for scallop photographs, Benjamin Grewe for providing data acquisition hardware, Ulrike Fuchs for optical design discussions, Hansjörg Kasper and Martin Wieckhorst for technical assistance, and Sascha Weidner for the design and production of 3D-printed parts. F.F.V. is supported by a HFSP fellowship (LT00687). T.N. received funding from H2020 Marie Skłodowska-Curie Actions (xenCAKUT - 891127). A.R. and S.H. were supported by a Dutch science foundation (NWO) VIDI Grant (#14637) and A.R. was additionally supported by an ERC Starting Grant (MULTICONNECT, #639938). In addition, this work was supported by grants from the Swiss National Science Foundation (grant nos. 31003B-170269, 310030_192617, and CRSII5-18O316 to F.H, and 310030_189102 to S.S.L.), European Union’s Horizon 2020 research and innovation programme to SSL (grant agreement No 804474, DiRECT), and the US Brain Initiative (1U01NS090475-01, FH).

## Contributions

F.F.V. and F.H. designed the project. F.F.V designed the objective, built the microscope, imaged samples, and analyzed data. A.M.R., T.N., S.H., M.S., A.L.H., A.B., S.C.F.N., A.R., E.T.S., S.S.L., and A.A. provided samples. L.W. created animations. F.F.V. and F.H. wrote the manuscript with input from all coauthors.

## Methods

### Objective design

The objective was designed using Zemax 2010 and Zemax OpticStudio 18. The aspherical correction plate was manufactured according to custom specifications out of fused silica by Asphericon Jena. The spherical mirror with 30-mm diameter was manufactured out of fused silica (Corning 7980) and coated with Aluminium and a protective layer of SiO_2_ (Praezisionsoptik Gera). The design process is described in Supplementary Note 1. For imaging in liquid media, the objective was filled with 65 ml of immersion medium.

### Setup

Excitation light from a tunable femtosecond Ti:Sapphire laser (Chameleon Ultra II, Coherent) was sent through a Pockels cell (Model 350-80 with 302RM controller, Conoptics) and expanded using a 6× Galilean telescope composed of a −25 mm and a 150 mm achromat (ACN127-25B and AC254-150B-ML, Thorlabs). The beam was then routed to a pair of 10-mm galvo mirrors (6220H, Cambridge Technology) and directed through a 4f-system composed of a f=89 mm scan lens (S4LFT0089/92, Sill Optics) and a f=300 mm tube lens (#88-597, Edmund Optics) to the objective. The galvos could be replaced by a resonant scanner / 8-mm galvo pair (CRS4K & 6220H, Cambridge Technology) for fast imaging. Emission light was separated from the excitation beam using a NIR/VIS dichroic (HC 705 LP, AHF) and collected onto two GAsP photomultipliers (H10771P-40 SEL, Hamamatsu) by a 4.7× demagnifying telescope composed of a f=90 mm achromat (G322389000, Qioptiq) and a wide-angle eyepiece (Panoptic 19 mm, TeleVue) as previously described^21^. Custom interchangeable filter cubes allowed the selection of emission channels. In addition, a 720-nm shortpass filter (ET720SP, AHF) located in front of the PMT blocked unwanted excitation light. PMT signals were amplified and lowpass-filtered by a transimpedance amplifier (DHPCA-100 set to 14-MHz bandwidth, 105 V/A gain, Femto). The amplified signal was digitized by a NI-5734 DAQ card connected to a NI-7961R FPGA (both National Instruments). Scanning waveforms were generated using NI-6341 cards in a PXIe-1073 chassis (National Instruments). ScanImage 2017b12 (Vidrio Technologies) running on MATLAB 2019b (MathWorks) was used to control image acquisition. The sample was moved in XYZ by a MP130-50-DC-L100 stage with linear encoders (Steinmayer Mechatronik) controlled by a Galil DMC-2132 motor controller. An additional rotation stage (PRM1Z8 with KDC101 controller, Thorlabs) allowed sample rotation. A second MP130 stage allowed XYZ translation of the spherical objective mirror. Both mirror and sample were quick-exchangeable using kinematic magnetic mounts (KB25/M, Thorlabs).

### PSF measurements

Sample holders for fluorescent beads were built by gluing smaller coverslips (3 mm ⌀, CS-3R-0, Warner Instruments) to a 2-mm glass capillary tube (Harvard Apparatus) using silicone adhesive (Wuerth Super RTV-Silikon). Fluoresbrite YG Microspheres (0.20 μm size; Polysciences Inc.) were diluted in distilled water and pipetted on the coverslip. The capillary tube was clamped into a sample holder and inserted into the Schmidt objective. For imaging at an index of n=1.45, the immersion chamber was filled with a fused-silica-matching liquid (50350, Cargille). Type A immersion oil (16482, Cargille) was used for measurements at n=1.51. The liquids were chosen such that the fluorescent beads did not dissolve; other immersion media such as BABB, DBE, and ECI rapidly dissolved Fluoresbrite beads. For each immersion medium and FOV location, 8 beads were measured and the resulting point spread functions (PSF) fitted with Gaussian profiles in XYZ.

### Imaging pollen grains

We glued a pollen granule sold as nutritional supplement (Morga Bluetenpollen, Morga AG) to a 2-mm borosilicate glass tube (TG200-4, Harvard Apparatus) with Super RTV Silicone Transparent (Wuerth). The dried granules were usually composed of a single type of pollen grain and could be preselected for shape and fluorescence under a widefield microscope. We selected a pollen granule with spikes (echinate sculptures). Immersing pollen granules in water or other polar solvents dissolves them into single pollen which is why we used ethyl cinnamate to image pollen granules in high refractive index medium in addition to imaging them in air.

### Functional imaging of larval zebrafish

We generated larvae by incrossing adult transgenic elavl3:GCaMP6s fish^22^. Small groups of ≈20 larvae were raised in filtered fish water in 10-cm Petri dishes on a 14h light, 10h dark cycle at constant 28°C. For imaging, larvae with strong GCaMP6s fluorescence and no pigmentation (*mitfa^−/−^*) were selected. Larvae were embedded in 2% agarose (UltraPure Low Melting Point Agarose, 16520-100, Invitrogen) before imaging. This set of animal experiments and procedures were performed in accordance with standard ethical guidelines and were approved by the Cantonal Veterinary Office of the Canton of Zurich.

### iDISCO staining and clearing of mouse brains

Mouse experiments and procedures were performed in accordance with standard ethical guidelines and were approved by the Cantonal Veterinary Office of the Canton of Zurich. Mouse brain chunks were stained for neurons and cleared using a modified version of the iDISCO protocol^3^. C57BL6/J wildtype and Thy1-YFP HJrs/J transgenic (Jackson, 003782) mice were anesthetized, perfused, and the brains post-fixed in 4% PFA in PBS for 4.5 hours at 4°C, shaking at 40 rpm. Mouse brains were washed in PBS for 3 days at RT and 40 rpm, with daily solution exchange. Samples were cut into 5 mm^3^ chunks and dehydrated in serial incubations of 20%, 40%, 60%, 80% methanol (MeOH) in ddH_2_O, followed by 2 times 100% MeOH, each for 1 hour at RT and 40 rpm. Pre-clearing was performed in 33% MeOH in dichloromethane (DCM) overnight (o.n.) at RT and 40 rpm. After 2 times washing in 100% MeOH each for 1 hour at RT and then 4°C at 40 rpm, bleaching was performed in 5% hydrogen peroxide in MeOH for 20 hours at 4°C and 40 rpm. Samples were rehydrated in serial incubations of 80%, 60%, 40%, and 20% MeOH in in ddH_2_O, followed by PBS, each for 1 hour at RT and 40 rpm. Permeabilization was performed by incubating the samples 2 times in 0.2% TritonX-100 in PBS each for 1 hour at RT and 40 rpm, followed by incubation in 0.2% TritonX-100 + 10% dimethyl sulfoxide (DMSO) + 2.3% glycine + 0.1% sodium azide (NaN3) in PBS for 5 days at 37°C and 65 rpm. Blocking was performed in 0.2% Tween-20 + 0.1% heparine (10 mg/ml) + 5% DMSO + 6% donkey serum in PBS for 2 days at 37°C and 65 rpm. Samples were stained gradually with polyclonal rabbit-anti-tyrosine hydroxylase antibody (Sigma Aldrich, AB152) 1:200 or polyclonal chicken-anti-GFP antibody (Aves Labs, GFP-1020) 1:400, followed by secondary donkey-anti-rabbit-AlexaFluor594 antibody (ThermoFisher, A32754) 1:200 or donkey-anti-chicken-AlexaFluor594 antibody (Jackson Immuno Research, 703-585-155) 1:400 in 0.2% Tween-20 + 0.1% heparine + 5% DMSO + 0.1% NaN3 in PBS (staining buffer) in a total volume of 1.5 ml per sample every week for 2 weeks at 37°C and 65 rpm. Washing steps were performed in staining buffer 5 times each for 1 hour, and then for 2 days at RT and 40 rpm. Clearing was started by dehydrating the samples in serial MeOH incubations as described above. Delipidation was performed in 33% MeOH in DCM o.n. at RT and 40 rpm, followed by 2 times 100% DCM each for 15 minutes at RT and 40 rpm. Refractive index (RI) matching was achieved in dibenzyl ether (DBE, RI = 1.56) for 4 hours at RT.

### Staining and clearing of *Xenopus* embryos

Whole-mount immunofluorescence procedures were adapted from previously described protocols^23,24^. All experiments were performed in accordance with standard ethical guidelines and were approved by the Cantonal Veterinary Office of the Canton of Zurich. Embryos were fixed at Nieuwkoop/Faber stage 42 for 40 minutes at room temperature in 4% PFA. Embryos were rinsed 3x with 1x PBS, dehydrated to 100% MeOH and stored overnight at −20C. Bleaching was performed at room temperature shaking under strong light in 10% H2O2 / 23% H2O / 66% MeOH for 48 hours. Embryos were rehydrated to 1x PBS with 0.1% Triton X-100 (PBT) and blocking was performed for 2h in 10% CAS-Block / 90% PBT (Life Technologies). Staining was performed using the Atp1a1 antibody (1:200, DSHB, A5) diluted in 100% CAS-Block for 48 hours at 4C. For nuclear counterstaining, DAPI (20 μg/ml, ThermoFisher, D1306) was added to the primary antibody mixture. Embryos were washed for 3x 30 min with PBT, blocked again for 2h (10% CAS-Block / 90% PBT) and incubated overnight at 4°C with secondary antibody (1:250, Alexa-Fluor-594, ThermoFisher A32742) diluted in 100% CAS-Block. Embryos were washed for 2x 1h with PBT and then 1h in PBS. Embryos were embedded in 2% low-melting agarose and dehydrated as follows: 75% MeOH/25% 1x PBS (2 hours), 50% MeOH/50% 1x PBS (2 hours), 25% MeOH/75% 1x PBS (2 hours), three times 100% MeOH (2x 45 min, 1x overnight). Clearing was performed in BABB (benzyl alcohol:benzyl benzoate 1:2) overnight.

### Neurofilament staining of a whole-mount chicken embryo

The embryo was sacrificed at day 4 of development and incubated in 4% paraformaldehyde for two hours at room temperature. For best staining and clearing results, the embryo was kept in constant, gentle motion throughout the staining procedure. Incubation was at 4°C. The tissue was permeabilized in 0.5% Triton X-100/PBS for one hour, followed by an incubation in 20 mM lysine in 0.1 M sodium phosphate, pH 7.3 for one hour. In a next step, the embryo was rinsed with five changes of PBS. Non-specific binding was blocked using 10% FCS (fetal calf serum) in PBS for one hour. The primary antibody mouse anti-neurofilament (1:2’000, RMO270, Invitrogen 13-0700) was added for 48 hours. The primary antibody was removed and the tissue rinsed with ten changes of PBS and an additional incubation overnight. After re-blocking in FCS/PBS for one hour, the embryo was incubated with the secondary antibody goat anti-mouse IgG-Alexa 568 (1:1’000, Jackson ImmunoResearch 115-165-003) for 15 hours. Then, the embryo was washed ten times with PBS followed by incubation overnight in PBS. For imaging the tissue was dehydrated in a methanol gradient (25%, 50%, 75% in ddH_2_O and 2x 100%, one hour each step) and cleared using 1:2 benzyl alcohol: benzyl benzoate (BABB) solution overnight (again gentle shaking is recommended for dehydration and clearing). The tissue and staining are stable for months when kept at 4°C in the dark. This set of animal experiments and procedures were performed in accordance with standard ethical guidelines and were approved by the Cantonal Veterinary Office of the Canton of Zurich.

### Human brain tissue preparation

The human occipital lobe samples were obtained from the body donation program of the Department of Anatomy and Embryology, Maastricht University. The tissue donor gave their informed and written consent to the donation of their body for teaching and research purposes as regulated by the Dutch law for the use of human remains for scientific research and education (“Wet op de Lijkbezorging”). Accordingly, a handwritten and signed codicil from the donor posed when still alive and well, is kept at the Department of Anatomy and Embryology Faculty of Health, Medicine and Life Sciences, Maastricht University, Maastricht, The Netherlands. The samples were obtained from a female patient (82 years) diagnosed with Alzheimer’s disease, dementia, aphasia, and depression. Brains were first fixed in situ by full body perfusion via the femoral artery. Under a pressure of 0.2 bar the body was perfused by 10 l fixation fluid (1.8 vol% formaldehyde, 20% ethanol, 8.4% glycerine in water) within 1.5–2 hours. Thereafter the body was preserved for at least 4 weeks for post-fixation submersed in the same fluid. Subsequently, brains were recovered by calvarial dissection and stored in 4% paraformaldehyde in 0.1 M phosphate buffered saline (PBS) until further processing. The samples were cleared and stained with MASH-AO as previously described^19^, with minor adjustments to the original protocol: The tissue was dehydrated in 20, 40, 60, 80, 100% methanol (MeOH) in distilled water for 1h each at room temperature (RT), followed by 1h in 100% MeOH and overnight bleaching in 5% H_2_O_2_ in MeOH at 4°C. Samples were then rehydrated in 80, 60, 40, 20% MeOH, permeabilized 2x for 1h in PBS containing 0.2% Triton X-100 (PBST). This was followed by a second bleaching step in freshly filtered 50% aqueous potassium disulfite solution. The samples were then thoroughly rinsed in distilled water 5x and washed for another 1h. For staining, the samples were incubated in 0.001% acridine orange solution in McIlvain buffer^25^ (phosphate-citrate buffer) at pH 4 for 5 days at room temperature. After 2.5 days the samples were flipped to allow for equal penetration of the dye from both sides. After staining, samples were washed 2x for 1h in the same buffer solution, dehydrated in 20, 40, 60, 80, 2x 100% MeOH for 1h each, and delipidated in 66% dichloromethane (DCM)/33% MeOH overnight. These steps were followed by 2x 1h washes in 100% DCM and immersion in ethyl cinnamate (ECi)^10^.

---

**Extended Data Fig. 1.**
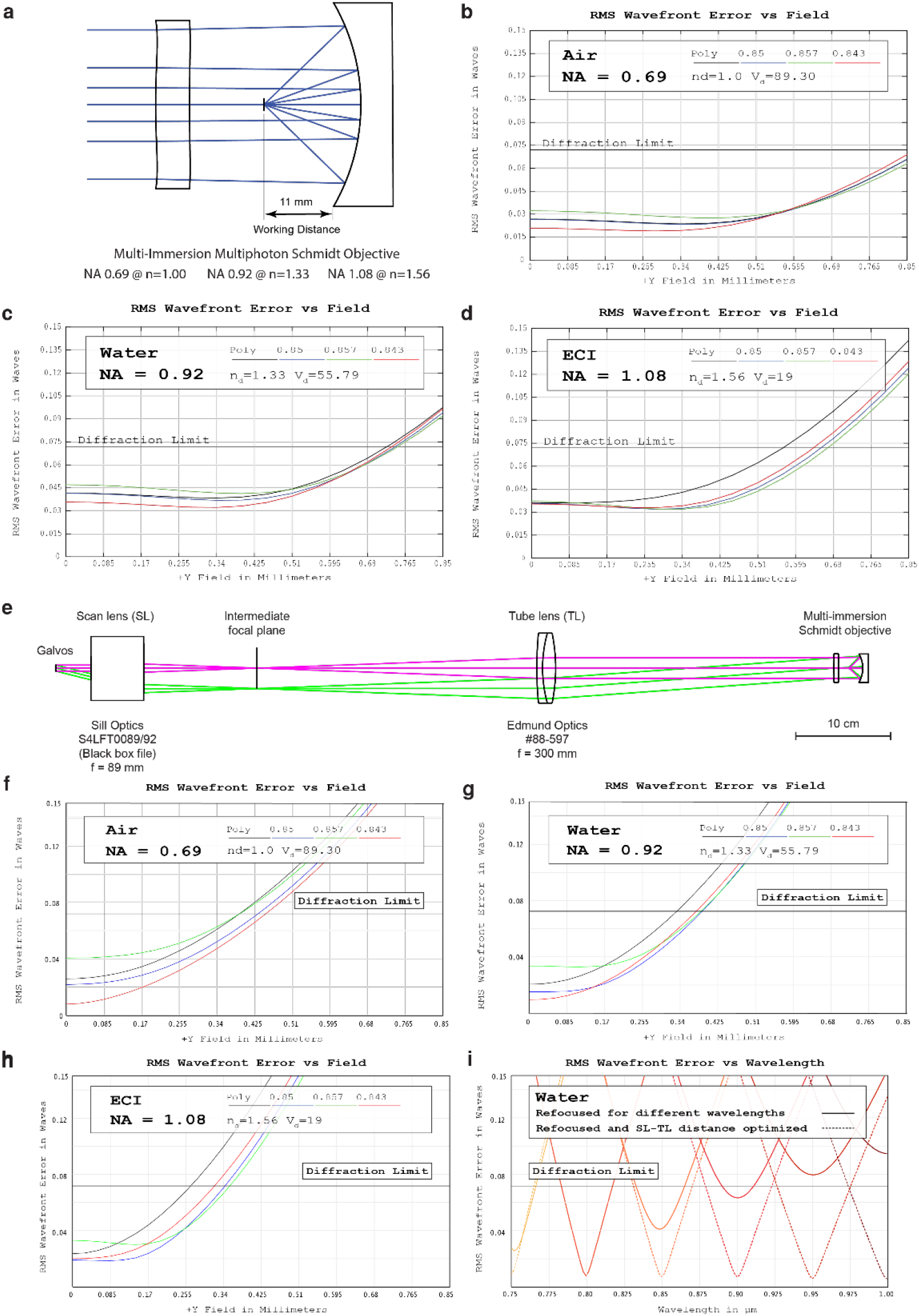
Theoretical performance of the Schmidt objective and the microscope: **a)** Cross-section of the optical design. **b)-d)** Root-mean-square (RMS) wavefront aberration vs. FOV of the objective in air, water, ECI, respectively. **e)** Cross-section of the laser-scanning path. **f)-h)** Root-mean-square (RMS) wavefront aberration vs. FOV of the objective in air plus scan path, water, ECI, respectively. Due to off-axis aberrations, the usable diffraction-limited FOV is reduced compared to the objective as a stand-alone item. **i)** On-axis RMS wavefront aberrations vs. wavelength. The system can be refocused to different wavelengths in the 750-1000 nm region. For best results, both the mirror-correction plate distance and the spacing of scan and tube lens can be optimized.

**Extended Data Fig. 2.**
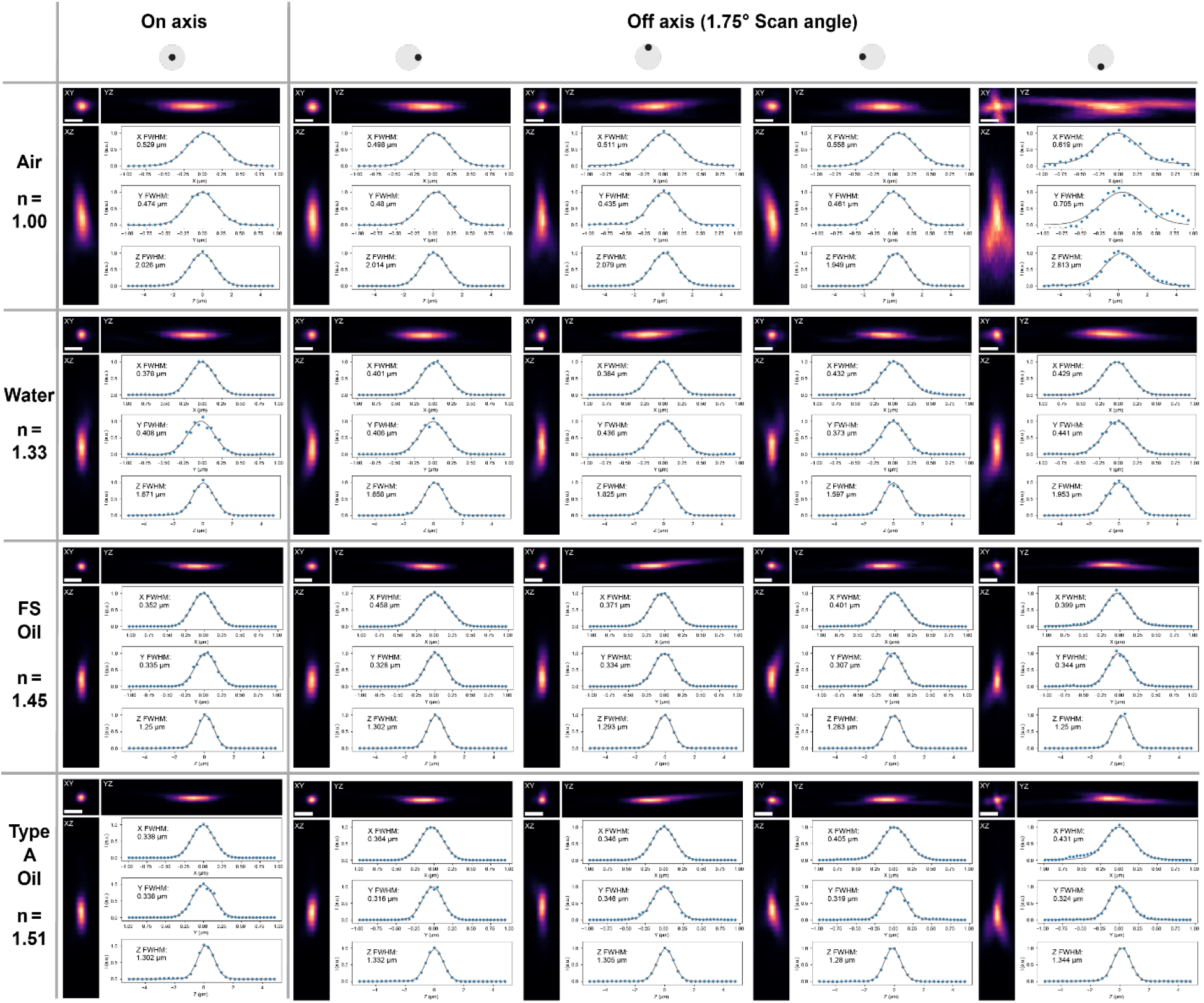
Example PSFs across the FOV in various media. Shown are XYZ maximum projections of example PSFs taken at different FOV positions in different immersion media ranging from air and water to fused silica matching oil (FS Oil, Cargille 50350), and Type A oil (Cargille 16482). In general, the PSF size decreases with index as the increased NA leads to higher resolution. However, beyond n=1.45, there is no change in resolution. We attribute this to residual aberrations from the minimally refractive inner surface of the correction plate: As the correction plate is made of fused silica with n=1.45, manufacturing-related surface irregularities have no effect at this index.

**Extended Data Fig. 3.**
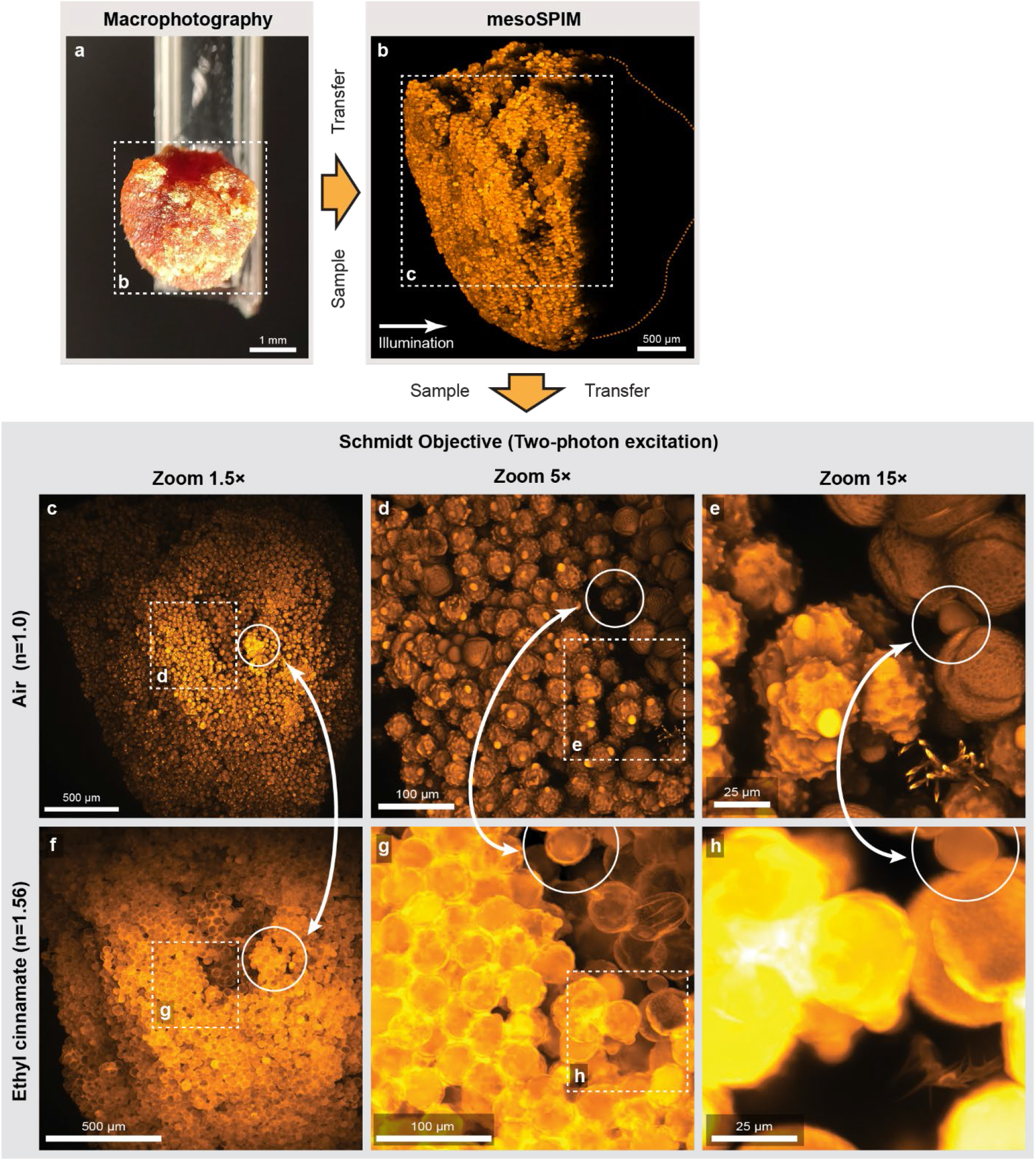
Imaging the same sample in air and in ethyl cinnamate (ECI). **a)** A pollen pellet composed of many individual pollen grains. **b)** Maximum projection of a single-photon light-sheet (mesoSPIM) stack of the sample. As the sample was illuminated from the side, only part of the pollen pellet is visible. **c)** Maximum projection of a stack taken in air after transferring the sample into the Schmidt objective. **d)-e)** Maximum projections of stacks taken at higher zoom. **f)** The immersion chamber was then filled with ethyl cinnamate (ECI) with an index of 1.56. This has several consequences: Firstly, at the same zoom level, the FOV shrinks by 1.56× as in the Schmidt objective, the product NA × FOV is constant. Secondly, the pollen are now much better index-matched and the microscope can also start visualizing the interior of the pollen grains. Therefore, the visual appearance of pollen grains is different than in air. **g)-h)** Zoomed-in maximum projections. Similar sample regions in both images are highlighted.

**Extended Data Fig. 4.**
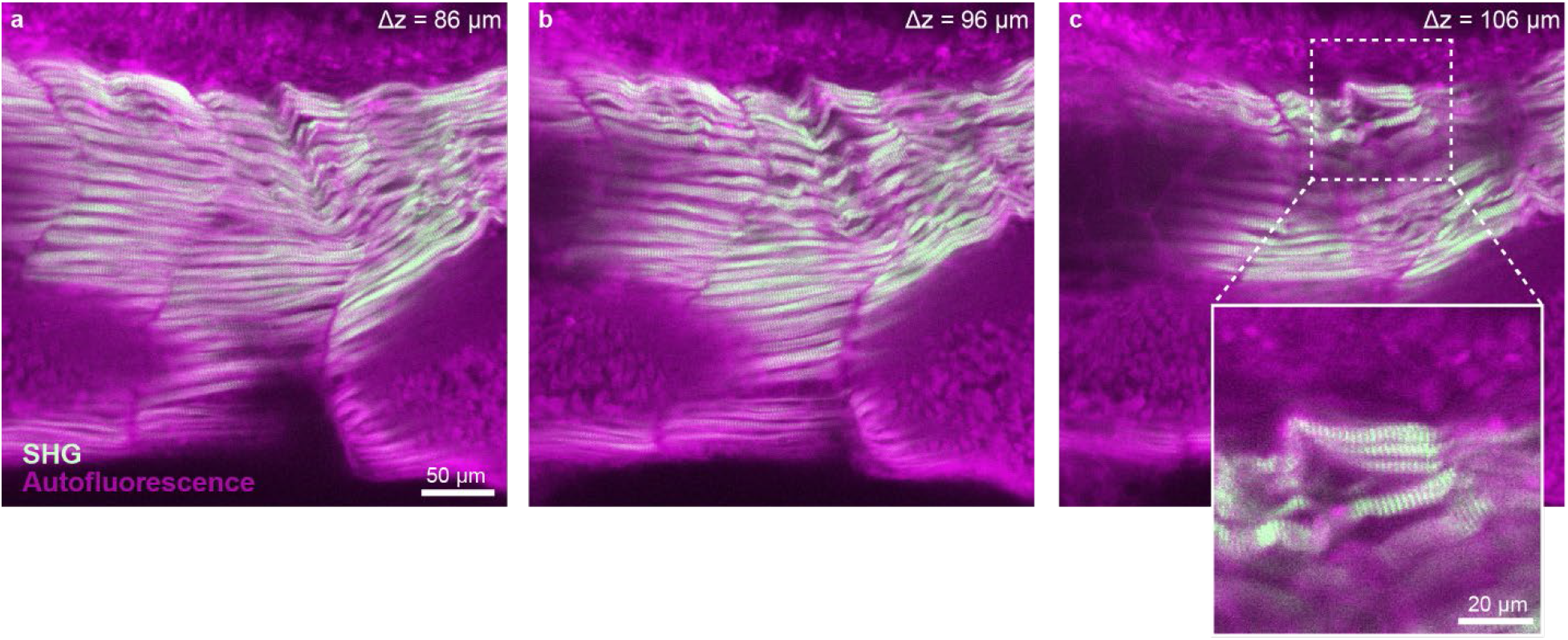
SHG imaging with the Schmidt objective. An unstained Xenopus embryo was embedded in Agarose and cleared using BABB. **a)-c)** Single z-planes from a stack taken in the the tail of the tadpole in the sagittal plane highlighting SHG signal from muscles

**Extended Data Fig. 5.**
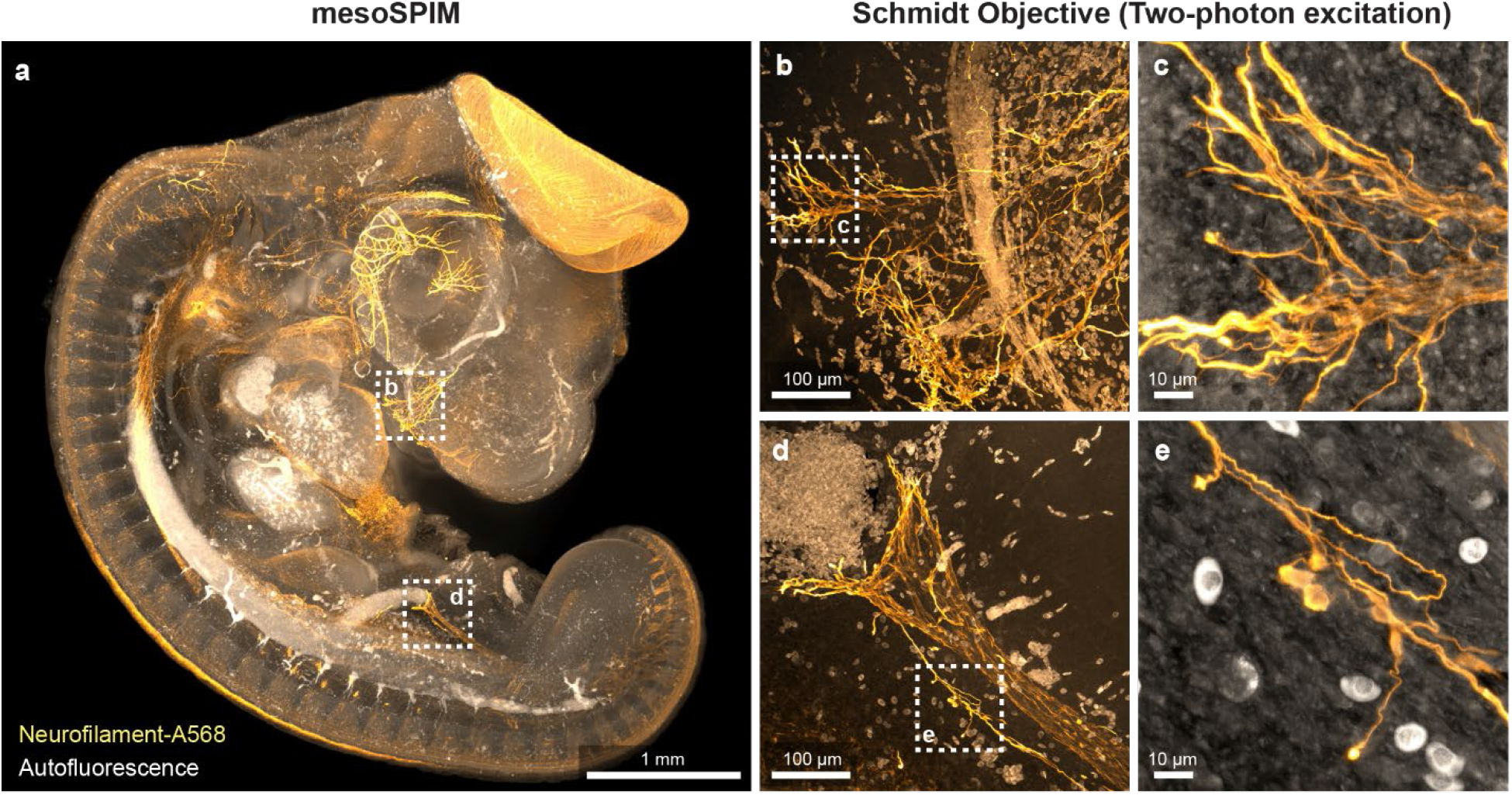
4-day Chicken embryo imaged with the Schmidt objective: A chicken embryo was stained for Neurofilament using Alexa 568 as a secondary antibody and cleared using BABB. **a)** Maximum projection of a single photon light-sheet (mesoSPIM) overview dataset of the sample. **b) & d)** Maximum projections of z-stacks taken at the highlighted locations in a) using the Schmidt objective and two-photon excitation. **c) & e)** Maximum projections of stacks taken at higher zoom level showing individual axons.

**Extended Data Fig. 6.**
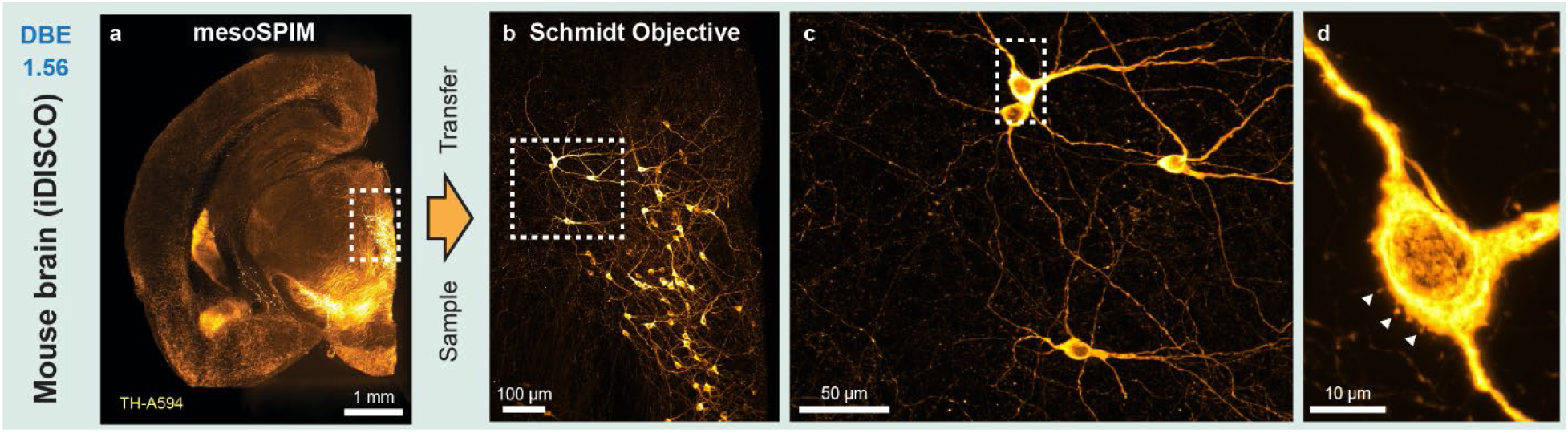
Dopaminergic neurons in the mouse brain imaged with the Schmidt objective: A 5 mm-thick coronal slice of a mouse brain was stained for Tyrosine Hydroxlase (TH) using Alexa 594 as a secondary antibody and cleared using Dibenyzl ether (DBE). **a)** Maximum projection of a single photon light-sheet (mesoSPIM) overview dataset of the sample. **b)** Maximum projection of a z-stacks taken at the highlighted location in a) using the Schmidt objective and two-photon excitation at 800 nm. **c) & d)** Maximum projections of stacks taken at higher zoom level showing individual dopaminergic neurons and spines on the cell body of a neuron (arrows).

**Extended Data Fig. 7.**
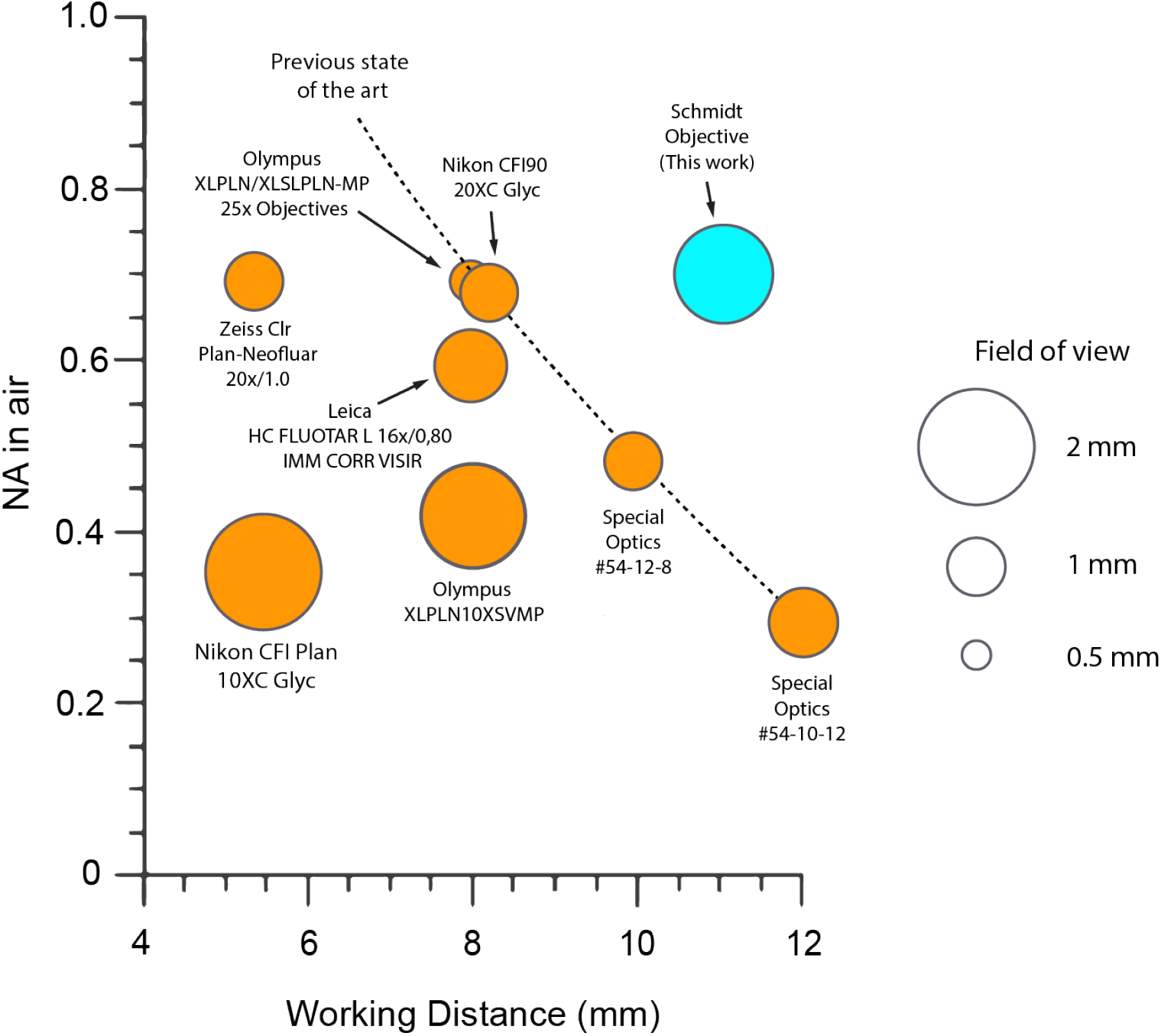
Comparison of multi-immersion microscope objectives for multiphoton imaging: In order to compare objectives with different index ranges, NA in air and working distance are used as a comparison metric. Only objectives with working distances larger than 4 mm are included in this comparison. The field of view is the theoretical FOV based on simulations or the field number provided by the manufacturers. Among the listed microscope objectives, the Schmidt objective design is the only one that will actually deliver a diffraction-limited image in air.

