## Supplemental Note 1 for "Reflective multi-immersion microscope objectives inspired by the Schmidt telescope"

### Supplemental Note 1: Schmidt Objective Theory

"After overcoming great difficulties (for this art has in reserve more difficulties than it seems to bear on its face), I at last succeeded in making the lenses which have provided me with the material for writing this account" (Christiaan Huygens, Systema Saturnium, 1655)

#### Introduction

In this supplementary note, we describe the theory of multi-immersion Schmidt microscope objectives from first principles. We begin by estimating the amount of spherical aberration generated by a spherical mirror, derive the surface equation for a Schmidt objective (or telescope) closely following the approach by Schroeder[1], and extend the formalism towards the multi-immersion case. Throughout our derivation we assume the paraxial and thin-lens approximations.

#### Spherical aberration of a spherical mirror

As a starting point for the design of a Schmidt objective, we consider the spherical aberration of a mirror:

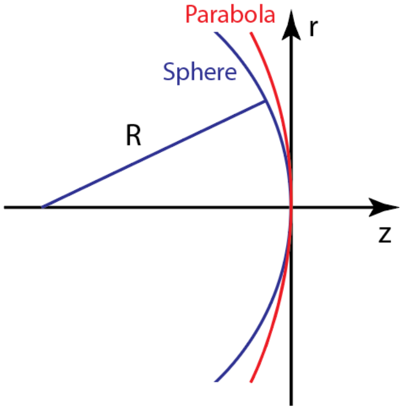


**Supplementary Figure 1.** Comparison of a circle and parabola. Both have their vertex located at the origin and have the same radius. We adhere to the common sign convention in optical design, namely that in the case shown, the radius R of the mirror is negative.

Here $r$ is the distance from the optical axis z. In Cartesian coordinates, $x^2 + y^2 = r^2$. A sphere with radius $R$ and with the vertex located at the origin has the equation:

$$R^2 = r^2 + (R - z_S(r))^2 \tag{1}$$

We then calculate $z_S(r)$ and neglect the case where the vertex is not located at the origin:

$$z_S(r) = R - \sqrt{R^2 - r^2} \tag{2}$$

This equation can be expanded into a Taylor series around = 0 :

$$z_S(r) = \frac{r^2}{2R} + \frac{r^4}{8R^3} + \frac{r^6}{16R^5} + \frac{5r^8}{128R^7} + \frac{7r^{10}}{256R^9} + \cdots \tag{3}$$

A parabola with radius $R$ would have the equation:

$$z_P(r) = \frac{r^2}{2R} \tag{4}$$

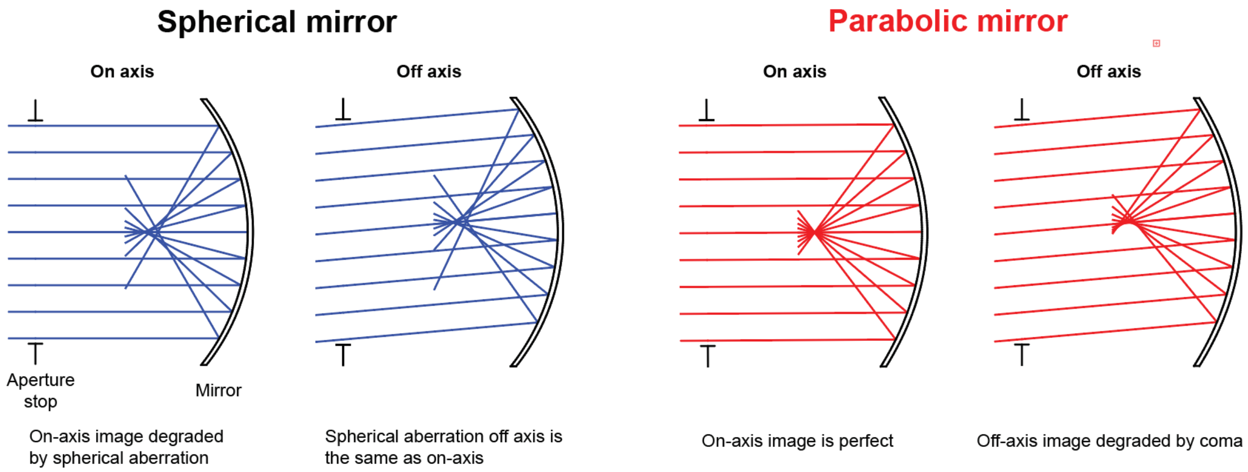


**Supplementary Figure 2.** Comparison of the on-axis and off-axis aberrations affecting a spherical and a parabolic mirror with the same radius of curvature. Here, the aperture stop is located at the center of curvature of the mirror. Left: The focus created by a spherical mirror suffers from spherical aberration: Rays hitting the mirror further away from the optical axis intersect it at a location closer to the mirror. However, the amount of spherical aberration is independent of the angle of incidence of the incoming bundle of rays. Right: A parabolic mirror creates a perfect focus on axis, however, off axis, the focus is degraded by coma.

For rays emanating from an on-axis object located at infinity, a parabola is known to bring all rays to a common focus free from aberrations. However, a spherical mirror will not bring all rays to a common focus: Supplementary Figure 2 shows that for a sphere and a parabola with identical $R$, the local curvature of the sphere is higher. Therefore, rays hitting the sphere at height $r$ are reflected with larger angular deviation compared to rays hitting the parabola at the same height. As a result, such rays fall short of the paraxial focus. This deviation from a perfect image is known as "spherical aberration" (Supplementary Figure 2). A parabolic mirror is, however, only capable of producing an aberration-free focus on the optical axis. The off-axis image is degraded by coma, an aberration that is named after the comet-like appearance of images of point sources (Supplementary Figure 2).

As the first term in Equations (3) and (4) is identical, their difference $\Delta z$ is proportional to the amount of spherical aberration generated by the mirror:

$$\Delta z = z_S(r) - z_P(r) = \frac{r^4}{8R^3} + \frac{r^6}{16R^5} + \frac{5r^8}{128R^7} + \frac{7r^{10}}{256R^9} + \cdots \tag{5}$$

To calculate the total spherical aberration, we need to estimate the optical path difference between two rays, one incident on the parabolic surface at height $r$ and one on the spherical surface at similar height. In the paraxial case, this difference is $2\Delta z$.

#### Design of a basic Schmidt objective

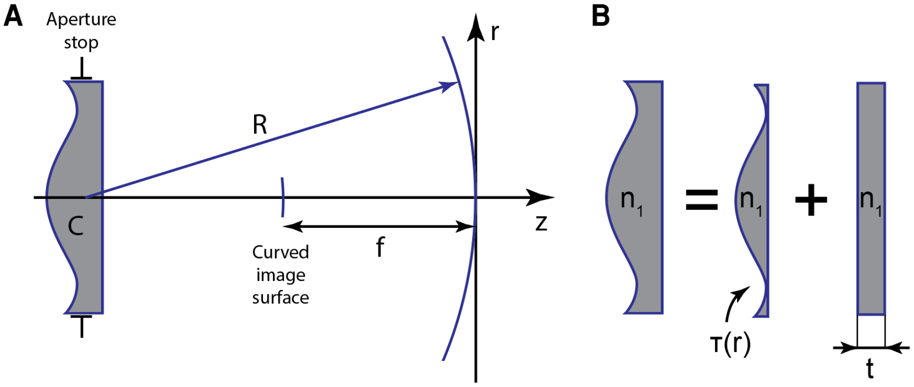


**Supplementary Figure 3. A)** Basic configuration of a Schmidt system. The center of curvature of the spherical mirror coincides with the aperture stop location. The correction plate C is located at this position as well. According to optical design convention, R is negative. **B)** The Schmidt plate is a glass plate with refractive index $n_1$ and consists of a plate with thickness t that does not introduce a radial optical path difference and a component with varying thickness $\tau(r)$.

A classical Schmidt objective[2] consists of three components (Supplementary Figure 3a): A spherical mirror, an aperture stop located at the center of curvature of the mirror and a correction plate at the stop location. Without correction plate, the system does not have an optical axis: As the curvature of the mirror is the same everywhere, rays from an off-axis object get reflected by a spherical surface of the same radius $R$. This means that there is no coma or astigmatism in this simple system and the only major aberration is spherical aberration. Due to the rotational symmetry of the system, the focal surface is curved with radius $R/2$ and the focal length is $f = R/2$.

The correction plate (or “Schmidt plate”) serves to compensate for the spherical aberration of the mirror. Compared to a parabolic mirror, the spherical mirror introduces a wavefront advance which needs to be compensated by an equal wavefront retardation by the corrector[1]. For terms up to $r^4$, the wavefront advance at the mirror is according to equation (5):

$$2\Delta z = \frac{r^4}{4R^3} \tag{6}$$

The corrector is a plane-parallel plate with thickness $t$ and index $n_1$. If we remove a layer of air of thickness $\tau$ at any height $r$ at the front of the correction plate and replace it with a layer of glass with optical path length $n_1\tau$, the net change in optical path length is $(n_1 - 1)\tau$ (Supplementary Figure 3b). As light propagation in glass with index $n > 1$ is slower than in air, this optical path difference is the required retardation if

$$(n_1 - 1)\tau = 2\Delta z = \frac{r^4}{4R^3} \tag{7}$$

Resolving for $\tau$, we find the required radial thickness profile

$$\tau(r) = \frac{r^4}{4(n_1 - 1)R^3} \tag{8}$$

Equation (8) defines the surface figure of an otherwise flat plate to correct the spherical aberration of the mirror. Here, we assume that this aspherical correction surface faces away from the mirror, but as only the sum of the wavefront retardation of the correction plate and wavefront advance by the mirror matter, it can also be facing the mirror.

However, while this type of surface figure compensates for spherical aberration, it also introduces an aberration known as spherochromatism or chromatic spherical aberration: Due to the dispersion $n_1(\lambda)$ of the glass material, third-order spherical aberration (equivalent to an optical path difference of 4$^{th}$ order) can only be compensated for a single wavelength. Other wavelengths will have varying residual spherical aberration. Therefore, it is advantageous to minimize spherochromatism by adding a weak additional lens shape (equivalent to a parabolic term) to the surface figure[1]. This additional lens will turn the resulting wavefront from a purely diverging shape into a combination of diverging and converging wavefronts and the correction plate into a combination of a positive and a negative lens (Supplementary Figure 4). As the dispersive effect of a lens inverts when changing its shape from positive to negative, the resulting "spectral spread" of the focus is reduced. Typically, this additional lens is chosen such that rays from a "neutral" zone around $r = \frac{\sqrt{3}}{2} r_0 \approx 0.866\, r_0$ (with $r_0$ as the maximum radius of the corrector) are undeviated[1]. In this case[1],

$$\tau(r) = -\frac{3{r_0}^2 r^2}{8(n_1 - 1)R^3} + \frac{r^4}{4(n_1 - 1)R^3} \tag{9}$$

This type of correction plate has its maximum thickness in the center (Supplementary Figure 4A) and acts like a convex lens in the center, like a plane-parallel plate around the neutral zone, and like a concave lens at large $r$ (Supplementary Figure 4B).

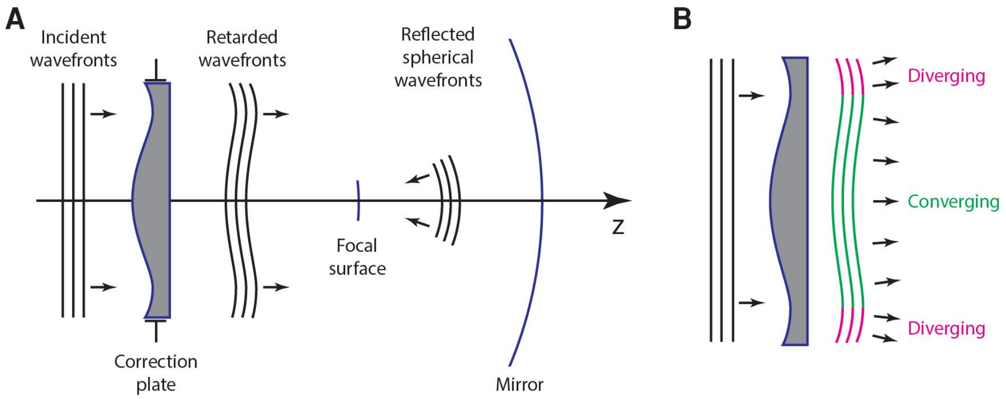


**Supplementary Figure 4. A)** Function of the correction plate. The incident parallel wavefronts are retarded by the corrector plate. This retardation counteracts the wavefront advance of the spherical mirror. As a result, the reflected wavefronts are spherical and converge towards the focus location. **B)** In the center, the correction plate acts like a convex lens whereas at the outer zones, it acts like a concave lens.

As we only included terms up to $r^4$, this type of surface can only correct third-order spherical aberration. At low focal ratios $f/D$ (with D being the aperture diameter $D = 2r_0$) or high numerical aperture ($NA = n \, sin \, \alpha$ ), spherical aberration of higher orders needs to be compensated as well. Therefore, $\tau(r)$ needs to be extended with higher-order terms. For microscope objectives with $NA \approx 1$, it is common that spherical aberration of 7$^{th}$ (and often 9$^{th}$) order needs to be corrected.

Please note that while the spherical mirror does not possess an optical axis, the correction plate does. As a result, off-axis image quality needs to be balanced against on-axis performance during optimization in an optical design program. Due to this requirement and the need for higher-order correction, the coefficients for the aspherical surface tend to deviate from Equation 9.

#### Design of a solid Schmidt objective

As a standard Schmidt telescope operates in air ("air-Schmidt"), the spherical mirror and correction plate are separated by an air gap with distance $d = 2f = R$. It is also possible to fill this gap with a solid or liquid medium with index $n_2$. If $n_1 = n_2$, such a design is referred to as a "solid-Schmidt" (Supplementary Figure 5A), which was commonly used for astronomical spectrographs in the mid-20$^{th}$ century[3,4]. To make such designs practical, a clamshell design was used (Supplementary Figure 5B).

Supplementary Figure 5A shows a solid-Schmidt with a chief ray entering the system at an angle $\theta$. Due to refraction, this ray propagates towards the mirror at an angle $\theta/n_1$. As the aperture stop coincides with the center of curvature, this ray is reflected back on itself. The height $h$ of the resulting focal surface is $f\theta/n_1$ from the z-axis. This means that the effective focal length of the solid-Schmidt is $f/n_1$ if $f$ is the focal length of an equivalent air-Schmidt. As a result, the focal ratio $f/D$ increases by a factor of $n_1$. Equivalently, the numerical aperture of the system increases by $n_1$. Thus, the image brightness (if used as a microscope objective) or "speed" (if used as a telescope) increases by ${n_1}^2$.

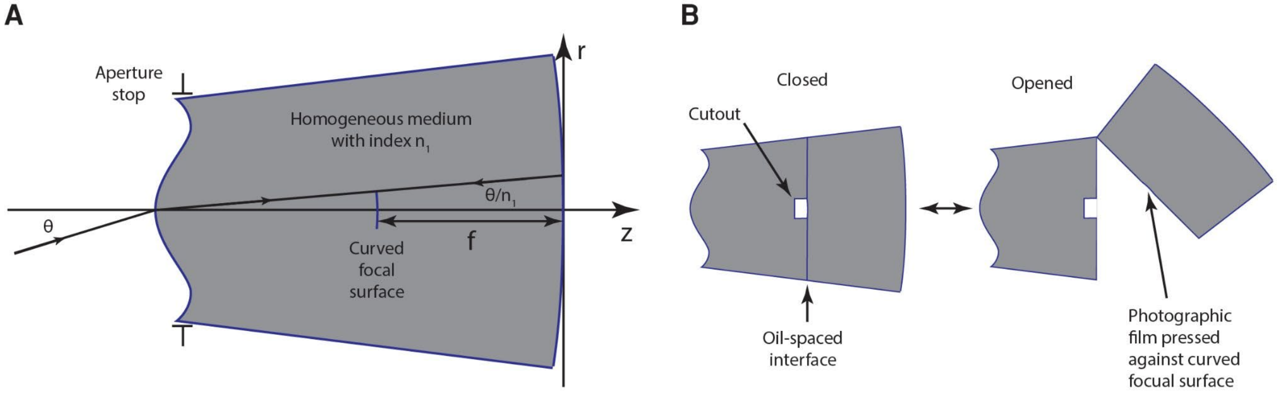


**Supplementary Figure 5. A)** Solid Schmidt with index $n_1$. A chief ray entering the system at an angle ϴ will propagate at an angle $\theta/n_1$ relative to the optical axis. **B)** Example clamshell design of a solid Schmidt utilized as a spectrograph camera.

When transitioning between two media with indices $n_i$ and $n_{i+1}$, a wavefront $w(r)$ is scaled by $(n_i/n_{i+1})\, w(r)$. Intuitively, the higher the index difference, the more the waveform is "flattened" after passing the interface. Thus, the surface figure $\tau_{solid}(r)$ of the correction plate for a solid-Schmidt needs to be $n_1$ times bigger than the equivalent air-Schmidt figure $\tau_{air}(r)$:

$$\tau_{solid}(r) = n_1\, \tau_{air}(r) = -\frac{3 n_1 {r_0}^2 r^2}{8(n_1-1)R^3} + \frac{n_1 r^4}{4(n_1-1)R^3} \quad (10)$$

As described above, this surface shape distorts the incoming parallel wavefront into a shape that counteracts the spherical aberration introduced by the spherical mirror. As no additional refraction occurs at the mirror (the law of reflection is independent of the refractive index: $\theta_1 = \theta_2$), the shape $w(r)$ of this corrected waveform inside the medium between mirror and correction plate has to be the same as for an air-Schmidt and thus:

$$w(r) = \frac{n_1-1}{n_1}\tau_{solid}(r) = -\frac{3{r_0}^2 r^2}{8R^3} + \frac{r^4}{4R^3} \quad (11)$$

#### Design of a multi-immersion Schmidt microscope objective

We now introduce a second aspherical surface into the system, which has a similar shape $w(r)$, according to Equation 11, as the corrected wavefront inside the system itself (Supplementary Figure 6). This surface has an interesting property: If we replace the medium between this surface and the mirror with a different one with index $n_2$, no refraction occurs at this surface: Because all passing rays are perpendicular to the new surface (according to the definition of a wavefront), they cannot get deviated by refraction, no matter what the difference in refractive index is. As no refraction happens, no additional aberrations are introduced. As a result, the mirror is able to focus the wavefront in a similar manner as in a solid Schmidt and the resulting image is corrected for spherical aberration independent of the refractive index $n_2$. This property is also approximately true for off-axis rays similar to the chief ray in Supplementary Figure 5 as long as $\theta$ is small. We therefore refer to a surface that is shaped similarly to the passing wavefront as a “minimally refractive surface”. By definition, such a surface does not have optical power and is thus of no use for correcting aberrations.

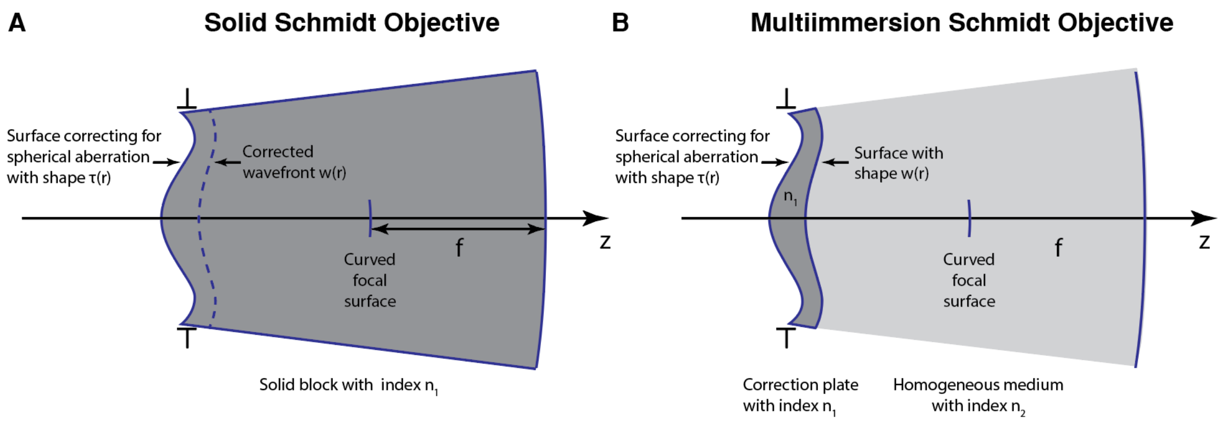


**Supplementary Figure 6**. Solid Schmidt vs. multi-immersion Schmidt objective. **A)** Cross-section of a Solid-Schmidt objective. The corrected wavefront w(r) inside the system is shown as a dashed line. **B)** Cross-section of a multi-immersion Schmidt objective. Here, a surface with shape w(r) is introduced as rear surface of the correction plate and the space between correction plate and mirror filled with a medium at index $n_2$. As all rays passing the rear surface are normal to the surface, no refraction occurs at this interface and no additional aberrations are introduced.

As the medium filling the space between correction plate and mirror can be a liquid, this type of Schmidt system constitutes a multi-immersion objective which is ideal for microscopy. If used as a microscope objective in combination with a laser scanning microscope, a sample is placed at the location of the focus and when scanning the beam, the laser focus will move along the curved focal surface. When used in combination with flat samples such as thin histological sections, only a ring-shaped zone will be in focus and a z-stack needs to be acquired to scan the entire sample. Thus, this type of objective is better suited for extended three-dimensional specimens, such as cleared samples, for which a curved imaging field is not a problem. As the system has only three optical surfaces – the two aspheres at the aperture stop and the spherical mirror, there are not enough degrees of freedom for correcting field curvature, which commonly requires a refractive corrector close to the image surface.

The properties of the multi-immersion Schmidt objective are comparable to the solid Schmidt system except that the bulk index $n_2$ of the medium can be arbitrarily modified:

- The numerical aperture of the objective is $NA = n_2 \sin\alpha$ . An increase in $n_2$ (by interchanging the medium) leads to an increase in NA by the same factor and a corresponding increase in lateral and axial resolution.
- The effective focal length of the system is $EFL = f/n_2$
- The size of the field-of-view is $FOV = FOV_{air}/n_2$
- If used as part of a compound microscope with magnification M in air, the effective magnification is $M_{eff} = M_{air}/n_2$
- The location of the focal surface is independent of $n_2$
- The curvature of the focal surface is approximately $R/2$ with $R$ being the radius of the mirror

In the case of a microscope objective operating at $NA \approx 1$, the shape of the front surface of a correction plate made from a medium of index $n$ typically requires terms up to 8th-order:

$$\tau_{highNA}(r) = a_2 r^2 + a_4 r^4 + a_6 r^6 + a_8 r^8 \quad (12)$$

The coefficients $a_i$ need to be found using numerical optimization in an optical design program. For a thin correction plate with index $n_1$, the shape of the required minimally refractive rear surface can then be found in analogy to Equation 6 as optical path differences are additive in the paraxial and thin-lens approximations:

$$w(r) = \frac{n_1 - 1}{n_1} \tau_{highNA}(r) \quad (13)$$

The dominant remaining aberration of the solid and multi-immersion Schmidt objectives is lateral chromatic aberration as shown in Supplementary Figure 7. Similar to Supplementary Figure 5, this figure shows a Schmidt system with a chief ray entering the system at an angle $\theta$. Due to dispersion, a short wavelength ("blue") ray is undergoing a larger angular deviation than a ray at longer wavelength ("red"). As a result, the image height increases with wavelength. If we assume that the thickness $t_C$ of the correction plate is small compared to the distance $t_{CM}$ between correction plate and mirror ($t_{CM} \gg t_C$), and that the liquid medium has a dispersion $n_2(\lambda)$, the height $h$ of the resulting image is $f\theta/n_2(\lambda)$. The difference in image height between two images at wavelengths $\lambda_1$ and $\lambda_2$ is thus:

$$\Delta h = \frac{f\theta}{n_2(\lambda_1)} - \frac{f\theta}{n_2(\lambda_2)} = f\theta\left(\frac{n_2(\lambda_2) - n_2(\lambda_1)}{n_2(\lambda_1) n_2(\lambda_2)}\right) \approx f\theta \frac{\Delta n_2}{{n_2}^2} \quad (14)$$

Thus, even if the combination of correction plate and mirror is optimized to be diffraction-limited, the system will not perform in a diffraction-limited fashion if the required wavelength range is chosen such that $\Delta h$ is larger than the lateral resolution of the objective $\Delta h \geq \Delta x = \lambda/(2NA)$.

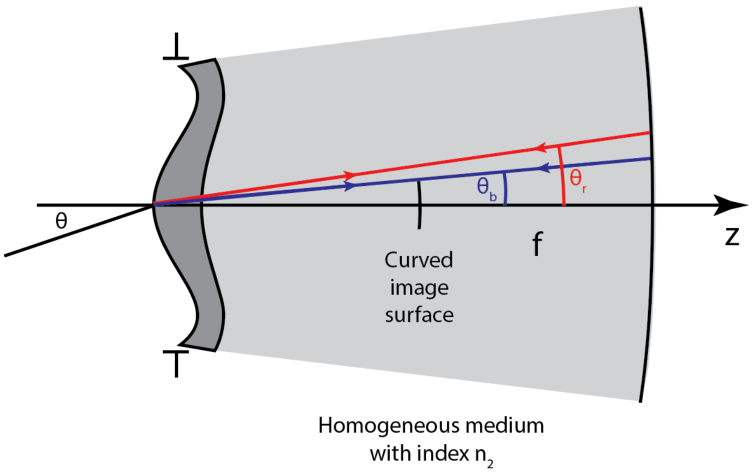


**Supplementary Figure 7**. Lateral chromatic aberration in a multi-immersion Schmidt objective: If the thickness of the correction plate is assumed to be much lower than the thickness of the liquid medium inside the objective, the dominant dispersive effect is solely due to the differential refraction of the chief ray. As blue rays are refracted less than red rays, the blue focus is formed closer to the optical axis compared to the red focus. If the required spectral bandwidth is small (i.e. when using multi-photon excitation with a single laser), all rays can be brought to a diffraction-limited focus.

If operation across a larger wavelength range is desired, lateral chromatic aberration needs to be corrected elsewhere in the optical system, for example by a custom tube lens. However, please note that the dispersion properties of liquids can vary dramatically – e.g ethyl cinnamate (ECI) has an Abbe number $V_D = 19.44$ whereas water has $V_D = 55.79$. Thus, the amount of lateral chromatic aberration introduced by such a custom tube lens needs to be tunable or a series of tube lenses needs to be constructed, each optimized for different media.

However, in the case of a two-photon microscope, the spectral width of commonly used ultrashort lasers is sufficiently small (approx. 10 nm for a 100-fs pulse at 850 nm) so that diffraction-limited imaging is possible if only a single excitation laser is used.

#### Design of a prototype multi-photon multi-immersion Schmidt microscope objective

Based on the paraxial design approach outlined above, we opted to realize a multi-photon Schmidt objective with the following parameters:

- Diffraction-limited imaging quality for n=1 (air) to n=1.56 (DBE or BABB)
- Diffraction-limited FOV diameter ≈ 1 mm
- NA > 1 for media with n=1.56, which means that NA > 0.64 in air

During system design, the index $n_1$ of the correction plate should be chosen to be in the middle of the desired index range for $n_2$ to minimize reflection losses at the interface. We chose fused silica with $n_D =$

1.4584 as its index is roughly in the middle between water $n_D = 1.33$ and ethyl cinnamate $n_D = 1.56$. In addition, fused silica is an ideal substrate for diamond turning of aspheres. We used Zemax 2010 and Zemax OpticStudio 21 to optimize the aspherical parameters of the correction plate. The mirror radius was chosen to be $r =$ -30.031 mm based on the test glass list of the manufacturer (POG Gera, Germany). The resulting design is shown in Supplementary Figure 8 and the design parameters are listed in Table 1. The aspherical correction plate was produced by Asphericon, Jena.

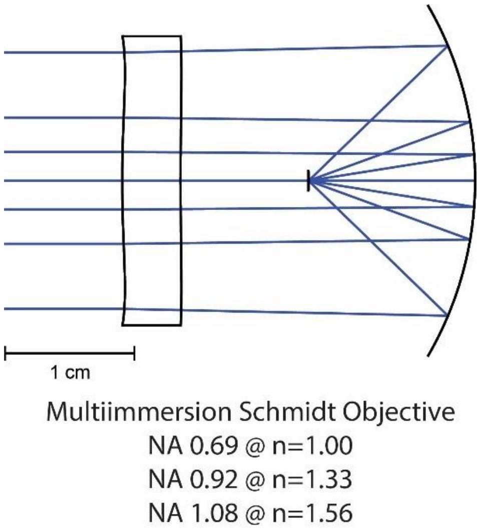


**Supplementary Figure 8**: Overview of the multi-photon multi-immersion Schmidt objective.

| Surface | Radius | Thickness | Material | $a_2$ | $a_4$ | $a_6$ | $a_8$ |
|---|---|---|---|---|---|---|---|
| 1 (Aperture Stop) | ∞ | 5 | Fused Silica | 5.173E-3 | -2.459E-5 | -2.768E-8 | -1.546E-10 |
| 2 | ∞ | 25.209 | Variable Immersion | 1.645E-3 | -8.564E-6 | -1.132E-9 | -6.447E-11 |
| 3 | -30.031 | -14.252 | Mirror | | | | |
| Image | -15.767 | | | | | | |

**Supplementary Table 1**. Design parameters of the prototype multi-photon multi-immersion microscope. Radii and thickness values are given in mm according to the sign convention in Supplementary Figure 1. The aperture stop is located at the first surface.

Extended Data Figure 1b-d shows the root-mean-square (RMS) wavefront aberration curves versus field size. For two-photon imaging at 850 nm with 100-fs pulses (typical values for a Titanium:Sapphire laser), the FWHM of the spectrum is approximately 14 nm. The system was designed for a Gaussian excitation beam with 22 mm $1/e^2$-diameter (apodization set to 1 in Zemax, aperture set to 22 mm). The aperture stop was set to coincide with the front surface of the correction plate (Surface 1). This means that – unlike in most microscope objective designs – the location of the aperture stop and back focal plane are the same.

After optimization of the aspherical parameters up to $8^{th}$ order, the design delivers a diffraction-limited FOV of D = 1.7 mm diameter in air at an NA of 0.69 (Extended Data Figure 1b). In water, the FOV shrinks to D = 1.4 mm diameter and the NA increases to 0.92 (Extended Data Figure 1c). In ECI, the NA reaches 1.08 and the FOV is D = 1.1 mm (Extended Data Figure 1d). The position and size of the aperture stop does not depend on the immersion medium. This means that the etendue (or optical throughput) of the optical system $G = \frac{\pi}{4} NA^2 D^2$ is constant. For our system $G \approx 1.3$ mm$^2$ , which is larger than the etendue of typical commercial multi-immersion objectives for cleared tissue such as the Olympus XLPLN10XSVMP that reaches G = 0.92 with a field number of 18 mm (10x magnification or D = 1.8 mm) and an NA of 0.6. Notably, commercial objectives are usually not diffraction-limited across the full FOV whereas our design is. One of the reasons why we can achieve this level of performance is that we allow the image surface to be curved with a radius of -15.767 mm. This means that for a FOV with 400 µm diameter, the sag of the image surface is 1.26 µm. This equals the change in Z-position between on-axis and off-axis regions in the image and is comparable to the size of the z-PSF. For a FOV with 1.1 mm diameter, the sag reaches 9.59 µm.

The other reason why it is possible to achieve a larger diffraction-limited FOV than commercial objectives with only three surfaces is that we optimized the objective for multi-photon imaging and realized that full color correction is unnecessary as long as the excitation pulses are sufficiently long (>70 fs) and only a single excitation wavelength is used at a time. This means that our system is essentially a "chromat". While it can be used with excitation wavelengths between 750 and 955 nm, two wavelengths with a separation wider than ≈30 nm will not be focused into a single diffraction-limited spot. Changing the wavelength of the excitation laser means that the system "refocuses" automatically to a different distance between focus and spherical mirror. In general, color correction for multi-immersion objectives with long working distances is challenging because not only the bulk refractive index of the liquid media can vary (typically $1.33 < n_d < 1.56$) but dispersion as well (ranging from $V_d$=19 for ECI to $V_d$=55 for water). This is part of the reason why the previously mentioned Olympus XLPLN10XSVMP objective (or other objectives from the Olympus lineup such as the XLSLPLN25XSVMP2 or the XLSLPLN25XGMP) is not recommended for widefield, confocal, or light-sheet usage as it has residual color aberrations. As outlined above, the dominant aberration in our design is lateral chromatic aberration, which means that the further the beam is scanned off-axis, the more the individual wavelengths of the excitation spot fall onto different radial locations.

#### Design of the excitation path

To build a cost-efficient two-photon microscope around our custom multi-immersion objective, we used off-the-shelf components for the scan and tube lens (Extended Data Figure 1e and Extended Data Figure 2a). To achieve the optimum resolution and large FOV, we needed to both overfill the back aperture and achieve a large scan angle at the aperture stop which in turn necessitates using large scan mirrors. We selected two swappable scan engines – either a galvo-galvo combination of two Cambridge Technology 6220H galvos with 10-mm mirrors or a resonant-galvo combination with a Cambridge Technology CRS-4k resonant scanner and a single 6220H galvo for the Y axis. For the scan and tube lens, we choose a Sill Optics S4LFT0089/92 (f=89 mm) and a f=300 mm achromat (#88-597, Edmund Optics) with 75 mm diameter. The excitation source was a Coherent Chameleon HP Ti:Sapphire laser. Prior to entering the scan engines, the beam was expanded using a 6× Galilean telescope composed of two achromats (-25 mm and 150 mm focal length; ACN127-25B and AC254-150B-ML, Thorlabs).

The combination of overfilling, large desired FOV, and restriction to off-the-shelf components results in a reduction of diffraction-limited FOV compared to a fully custom scan path. For example, in water, the simulated diffraction-limited FOV shrinks to a diameter of 680 µm (Extended Data Figure 1g) compared to 1.4 mm for the stand-alone design (Figure 5D). This is largely due to the introduction of additional lateral chromatic aberration by the scan lens/tube lens combination. Despite this drastic reduction, the theoretical diffraction-limited FOV is still comparable with commercial objectives. For example, the Olympus XLSLPLN25XGMP 25x-objective is designed for a field number of 18 mm, which means that its maximum FOV is ≈720 µm. It can be assumed that this system is not diffraction-limited under similar excitation conditions as the multi-immersion Schmidt objective (two-photon excitation at 800 nm with 100 fs pulses).

#### Alignment of the objective

Apart from imaging performance, the optical design of a microscope objective also has to fulfill tolerancing and alignment specifications. In high-NA microscope objectives, lens spacing, decentration, tip/tilt, and glass tolerances are usually very tight, which means that specialized assembly methods such as alignment turning are required. Alignment turning is a high-precision technique, in which a lens glued in a mount is centered on a specialized lathe by cutting away its housing such that the axes of revolution of mount and the optical axis match. Also, batch-to-batch variations in refractive index commonly require melt matching, the reoptimization of the optical design for a specific batch of glass. In this technologically highly demanding engineering space, our Schmidt objective occupies an outlier position in terms of manufacturing tolerances and alignment process: While the production of the bi-aspheric correction plate requires careful tolerancing, the actual assembly and alignment of the objective is extremely straightforward compared to standard high-NA objectives: Given that the optical axis of a Schmidt system is solely defined by the optical axis of the correction plate as the spherical mirror does not have an axis of symmetry, the only degrees of freedom are the XYZ translation of the mirror relative to the correction plate. Any tip/tilt of the mirror can be compensated by an offset in XYZ mirror position. Therefore, we opted to place the mirror on a motorized XYZ-stage as this would allow straightforward exchange of the mirror for cleaning in our prototype. In addition, the multi-immersion capability of the system means that it can be aligned in air and then filled with the immersion medium of choice. The alignment process itself is also rather simple: Any XY offset of the mirror relative to the optimum position results in a relatively uniform alignment coma over the full FOV. This is comparable to alignment coma in classical Schmidt telescopes[7] and can be minimized simply by canceling this on-axis coma by moving the mirror in XY. In practice, we use a test sample with 1-µm fluorescent beads deposited on a coverslip with 3 mm diameter and visually optimize the on-axis PSF. Typically, the mirror XY position needs to be optimized within ±10 µm. The same test sample can also be used to set the Z-distance between correction plate and mirror. In this case, the lateral size of the on-axis PSF does not indicate whether the distance is set correctly; in fact, deviations of ±200 µm are possible without any drastic degradation in spot size. On the other hand, off-axis PSFs exhibit coma when this distance is incorrect. This coma is minimal at the correct distance and changes from overcorrected to undercorrected when passing through the minimum. Taken together, this means that the entire alignment process is much simpler for the Schmidt objective compared to any other objective of comparable NA and FOV.

#### Estimating transmission and reflection losses

Because it is not possible to directly place a power meter sensor in the focus of the Schmidt objective, as it would obstruct the excitation beam, we need to estimate the transmission losses of the system based on the known properties of the media and coatings. The key wavelengths are 500 nm (emission), 800 nm (our most common wavelength for imaging in samples cleared with organic solvents), and 920 nm (wavelength used for imaging GCaMP6s in zebrafish larvae):

- The material of the correction plate is uncoated fused silica. Based on the Fresnel equations, we thus expect reflection losses of 3.5% (or 96.5% transmission) over this wavelength range at the first surface.
- The reflection losses at the interface between correction plate and liquid are negligible given the small index differences ($\Delta n \approx 0.12$ leading to reflectivities of 0.4% in water or 0.13% in ECI). When imaging in air, this loss has to be taken into account, though.

- For normal incidence and in air, the coating on the spherical mirror has a reflectivity of 90.9% at 500 nm, 79.5% at 800 nm, and 84.2% at 920 nm (Measurements by POG). We can estimate the wavelength dependence of the reflectivity $R$ for different media in contact with the mirror based on the Fresnel reflection from a metal at normal incidence:

$$R = \frac{(n_{metal}(\lambda)-n_{liquid}(\lambda))^2 + \kappa_{metal}{}^2}{(n_{metal}(\lambda)+n_{liquid}(\lambda))^2 + \kappa_{metal}{}^2} \quad (15)$$

  Here, $n_{metal}(\lambda)$ is the wavelength-dependent index of the metal and $n_{liquid}(\lambda)$ is the index of the liquid medium. $\kappa_{metal}$ is the extinction coefficient of the metal[5]. Based on this equation, we can predict that the reflectivity in water is approximately 3.5% less than in air (i.e. 87.7% at 500 nm, 76.7% at 800 nm, and 81.2% at 920 nm). For media with an index of 1.56 (DBE, BABB, and ECI), we predict that the reflectivity is approximately 7% less than in air (i.e. 84.5% at 500 nm, 74% at 800 nm, and 78.3% at 920 nm). For this estimate, the effect of the protective quartz coating on the mirror is neglected.
- The excitation beam has to pass through the liquid medium with a path length of ≈ 4 cm:
  - For water[6], this results in a transmission of 99.9% at 500 nm, 92.4% at 800 nm, and 59.7% at 920 nm.
  - With a Coherent Chameleon Ti:Sa laser tuned to 800 nm, we find a transmission of DBE through a 100-mm Hellma 100-OS cuvette (corrected for reflection losses at the cuvette windows) of 98 +/- 2%. For ECI and BABB, we measure 97 +/- 1% and 98 +/- 1%, respectively (mean +/- s.d., n = 3 measurements with a Thorlabs PM100D power meter with S425C sensor). We can thus estimate that for an optical pathlength of 4 cm inside the Schmidt objective, transmissions are likely >99%.

Taken together, we estimate the total transmission of the multi-immersion Schmidt objective to be:

- In air: 84% at 500 nm, 73% at 800 nm, and 78% at 920 nm
- In water: 85% at 500 nm, 68% at 800 nm, and 47% at 920 nm
- In media with n=1.56 (DBE, BABB and ECI): 81% at 500 nm, 70% at 800 nm, and 75% at 920 nm

While both the reflectivity of the mirror and the correction plate could be improved using better coatings, the most significant cause of transmission loss stems from water absorption at 920 nm. This is an effect not specific to the multi-immersion Schmidt objective. Any microscope objective used for multi-photon imaging (or any other NIR illumination) will show a significant transmission loss if working distances reach several centimeters. Heating of water due to bulk absorption is negligible, though. Neglecting any heat dissipation from the water to the chamber and with 200-mW excitation power (far exceeding the maximum values for our imaging examples) at the aperture stop at 920 nm, the 80 mW lost due to bulk absorption would lead to an increase in temperature of the 65 ml liquid volume by 1°C only after 56 minutes. Given that we use pulsed femtosecond excitation, it is likely that most of the heat dissipates. Thus, whereas our Schmidt objective shows low (<50%) transmission for imaging GCaMP in living zebrafish in sea water, the design operates at transmission levels >70% in typical organic solvents (without any optimization of coatings) and is thus comparable to commercial objectives. While it is clearly suboptimal, we specifically chose the Aluminum coating with protective $SiO_2$ layer by POG during the design process because it was not clear whether typical coatings would get damaged by the repeated immersion and cleaning cycles during routine use of the objective. DBE and BABB are usually considered to be very corrosive and often advised against when using commercial microscope objectives due to damage to glued lenses and possibly coating. To check for long-term degradation by either DBE or BABB, we immersed two test plates for 6 months in these media and checked whether the reflectivity changed. For this, we used a 647 nm laser beam (Omicron SOLE-6) expanded to a 10-mm spot and reflected off the mirror at an angle of 30 degrees (in air). We used a Thorlabs PM100D power meter with S121C sensor. Prior to immersion, we measured a reflectivity of 90.6% and 90.3%. After 6 months of immersion, the reflectivity change was minor: 88.5% (DBE mirror) and 88.4% (BABB mirror) and well within the typical +/-2% accuracy range of standard power meters. Both before and after long-term immersion, the values were close to the specifications of the coating (88.3% at 647 nm).
